# The ups and downs of amino acid co-evolution: evolutionary Stokes and anti-Stokes shifts

**DOI:** 10.1101/2020.08.31.271775

**Authors:** Noor Youssef, Edward Susko, Andrew J. Roger, Joseph P. Bielawski

## Abstract

The most fundamental form of epistasis occurs between residues within a protein. Epistatic interactions can have significant consequences for evolutionary dynamics. For example, a substitution to a deleterious amino acid may be compensated for by replacements at other sites which increase its propensity (a function of its average fitness) over time - this is the evolutionary Stokes shift. We discovered that an opposite trend -the decrease in amino acid propensity with time-can also occur via the same epistatic dynamics. We define this novel and pervasive phenomenon as the evolutionary anti-Stokes shift. Our extensive simulations of three natural proteins show that evolutionary Stokes and anti-Stokes shifts occur with similar frequencies and magnitudes across the protein. This high-lights that decreasing amino acid propensities, on their own, are not conclusive evidence of adaptive responses to a changing environment. We find that stabilizing substitutions are often permissive (*i.e*., expand potential evolutionary paths) whereas destabilizing substitutions are restrictive. We show how these dynamics explain the variations in amino acid propensities associated with both evolutionary shifts in propensities.

## Introduction

Amino acid interactions within a single protein are the most fundamental form of epistasis. Epistatic interactions between sites can occur because of functional, structural, or stability constraints (Ortlund *et al*., 2007; Pollock *et al*., 2012; Gong *et al*., 2013). Here we focus on the latter constraints on stability by using a model for protein evolution based on thermodynamic principles. This modeling framework has been shown to reproduce realistic evolutionary dynamics with regards to protein stability values (Goldstein, 2011), evolutionary rates (Youssef *et al*., 2020), and convergence rates (Goldstein *et al*., 2015). In this article, we explore the evolutionary dynamics that arise due to nonadaptive stability-constraints on proteins.

Under nonadaptive evolution, a protein evolves on a fixed fitness landscape with no changes in environment or function (Wright, 1932). Natural selection plays an important role in maintaining the protein near a peak on its fitness landscape. Evolutionary dynamics associated with a fitness peak reflect the equilibrium between mutation, drift and selection. At equilibrium, most mutations are deleterious while a small proportion may be beneficial. The higher probability of fixations of the fewer but more advantageous mutations is balanced by the lower probability of fixations of the more frequent yet disadvantageous mutations, resulting in an equal proportion of deleterious and beneficial substitutions (*i.e*., fixed mutations) at equilibrium (Goldstein, 2013; Cherry, 1998). This stands in sharp contrast to the expected dynamics under adaptive evolution, where a change in protein function or environment (and hence a change in the fitness landscape) renders the current state of the protein suboptimal for the new conditions. The shift in landscape is followed by successive fixations of beneficial substitutions as the protein adapts towards the new fitness peak (dos Reis, 2015; Jones *et al*., 2017).

Since its origin (Kimura, 1968), the strictly neutral model of protein evolution remains the most frequently used null scenario that must first be rejected in order to postulate a history of adaptive evolution (Kimura, 1968, 1991; Duret, 2008). Over the past decade, researchers have shown that equilibrium dynamics under more complex models, informed by the selective constraints for protein-stability, produce equilibrium dynamics that are largely consistent with neutral theory (*e.g*. with regards to the distribution of mutational fitness effects (Goldstein, 2011)). Using these nonadaptive stability-informed models, researchers have observed that various evolutionary phenomena characteristic of natural proteins can arise without the need for invoking adaptive evolution. For example, Goldstein (2011) used a thermodynamic model to argue that the marginal stability observed in many natural proteins can emerge from a simple balance between mutation, drift, and selection, challenging the widely held notion that evolution actively selects for marginal stability (DePristo *et al*., 2005).

Additionally, within the same modeling framework, Pollock *et al*. (2012) observed that the propensity for a resident amino acid at a site tends to increase over time due to compensatory substitutions at other sites in the protein. (In this context, propensity is the frequency of an amino acid arising at a site for a fixed background sequence.) They referred to this phenomenon as the evolutionary Stokes shift. Risso *et al*. (2014) found empirical evidence of evolutionary Stokes shifts in the evolution of thioredoxins and *β*-lactamases; however, those shifts in propensities were minor. Interestingly, Popova *et al*. (2019) recently observed the opposite trend where the fitness of the resident amino acids decreased with time - they termed this phenomenon “senescence”. Popova *et al*. (2019) contend that the decrease in preferences must be the result of adaptive protein evolution, and that, unlike the evolutionary Stokes shifts, mere epistatic constraints “cannot lead to a systematic reduction in fitness of the incumbent alleles”.

Do resident amino acid preferences tend to increase, decrease, or remain relatively conserved throughout a protein’s evolution? And to what extent are they shaped by adaptive or nonadaptive processes? Using extensive simulations under a thermodynamic model for protein stability, we show that all three trajectories emerge from the nonadaptive dynamics at mutation-drift-selection equilibrium. Importantly, we describe a novel phenomenon whereby resident amino acid preferences can decrease merely due to epistatic constraints – which we call the evolutionary anti-Stokes shift. We then show that evolutionary anti-Stokes shifts are as common as Stokes shifts, and characterize the underlying mechanisms that give rise to them. In line with experimental evidence (Gong *et al*., 2013) we found that stabilizing substitutions are permissive (*i.e*., expand potential evolutionary paths) whereas destabilizing substitutions are restrictive. We show how these dynamics explain the variations in amino acid propensities associated with evolutionary Stokes and anti-Stoke shifts.

## Results

We use a thermodynamic model of protein evolution where fitness is equal to the probability of observing an amino acid sequence in a native protein structure at equilibrium (which is a function of its stability, Δ*G*). We assume no changes in protein structure or function so that the global fitness landscape (*i.e* the mapping between amino acid sequences and fitness values) remains constant. Nonetheless, this modeling framework accounts for stability-mediated epistasis by permitting differences in the marginal site-specific fitness landscapes depending on the residues occupying other sites in the protein (i.e. the background protein sequence). Throughout the simulations, we calculate the site-specific fitness landscape 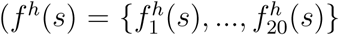 for a given site *h* and background sequence *s*), at all sites and given all observed sequences. Amino acids that confer higher fitness values (improve stability) will tend to more frequently occupy the site and will, therefore, have higher expected frequencies at equilibrium for a given background sequence. In this way, the frequency of an amino acid is related to its fitness. As a result, frequency landscapes are similarly site-specific and context-dependent 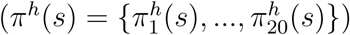. Note, however, that the fittest amino acid may not necessarily be the most frequently observed residue. This can occur when a suboptimal amino acid has many codon aliases - the high number of synonymous codons and/or mutational bias can increase the propensity for the residue despite having slightly lower fitness.

An evolutionary Stokes shift is a phenomenon whereby the propensity 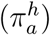 for the resident amino acid at that site increases over time due to compensatory substitutions at other positions in the protein (Pollock *et al*., 2012). The propensity for an amino acid is its equilibrium frequency given a fixed background protein sequence,

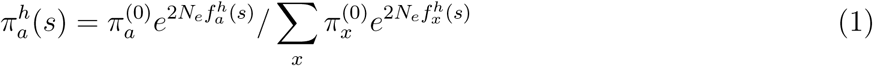

where *N*_*e*_ is the effective population size and 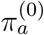 is the stationary frequency of amino acid *a* in the absence of selection pressure (dos Reis, 2015). In their formulation, Pollock *et al*. (2012) assume that 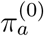 are uniform 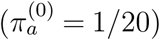. This would be true if all amino acids were specified by the same number of codons and there were no underlying mutational biases. However, our simulations are based on three proteins with DNA mutation parameters estimated from multiple sequence alignments of their natural codon sequences. Importantly, analyses of the three protein genes suggest the presence of mutational biases (with unequal nucleotide frequencies, and transition/transversion rate biases). We account for these mutational biases by estimating protein-specific 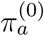 given the estimated mutation parameters for each protein (figure S1). The following results are based on 500 protein-specific simulations for three proteins with PDB structures 1qhw, 2ppn, and 1pek (see Methods). We ran each simulation for 500 substitutions.

### Both evolutionary Stokes and anti-Stokes shifts emerge from nonadaptive stability-constraints on protein fitness

Throughout our simulations, and in real protein evolution (Risso *et al*., 2014; Gong *et al*., 2013; Ashenberg *et al*., 2013), the propensity for certain amino acids relative to other amino acids changes over time. In natural proteins, these variations may be due to global constraints on protein stability, or are related to functional restrictions. By contrast, the fitness model we employ assumes selection acting only on protein stability. Therefore, any variations in sites’ fitness and propensity landscapes are solely due to stability-induced epistatic interactions between sites (and not due to external changes in environment or function). Examples of these propensity dynamics from a simulation of the 1pek protein are shown in figure 1. The propensity for aspartic acid (D), the resident amino acid at site 232, changes considerably as substitutions occur at other positions in the protein (figure 1A). In this case, the site experiences an evolutionary Stokes shift where its propensity increases over time due to compensatory substitutions at other positions (figure 1A). In contrast, within the same simulation, the propensity for proline (P), the resident amino acid at site 72, decreases as substitutions occur at other positions (figure 1B). We refer to this phenomenon as the evolutionary anti-Stokes shift.

**Figure 1:**
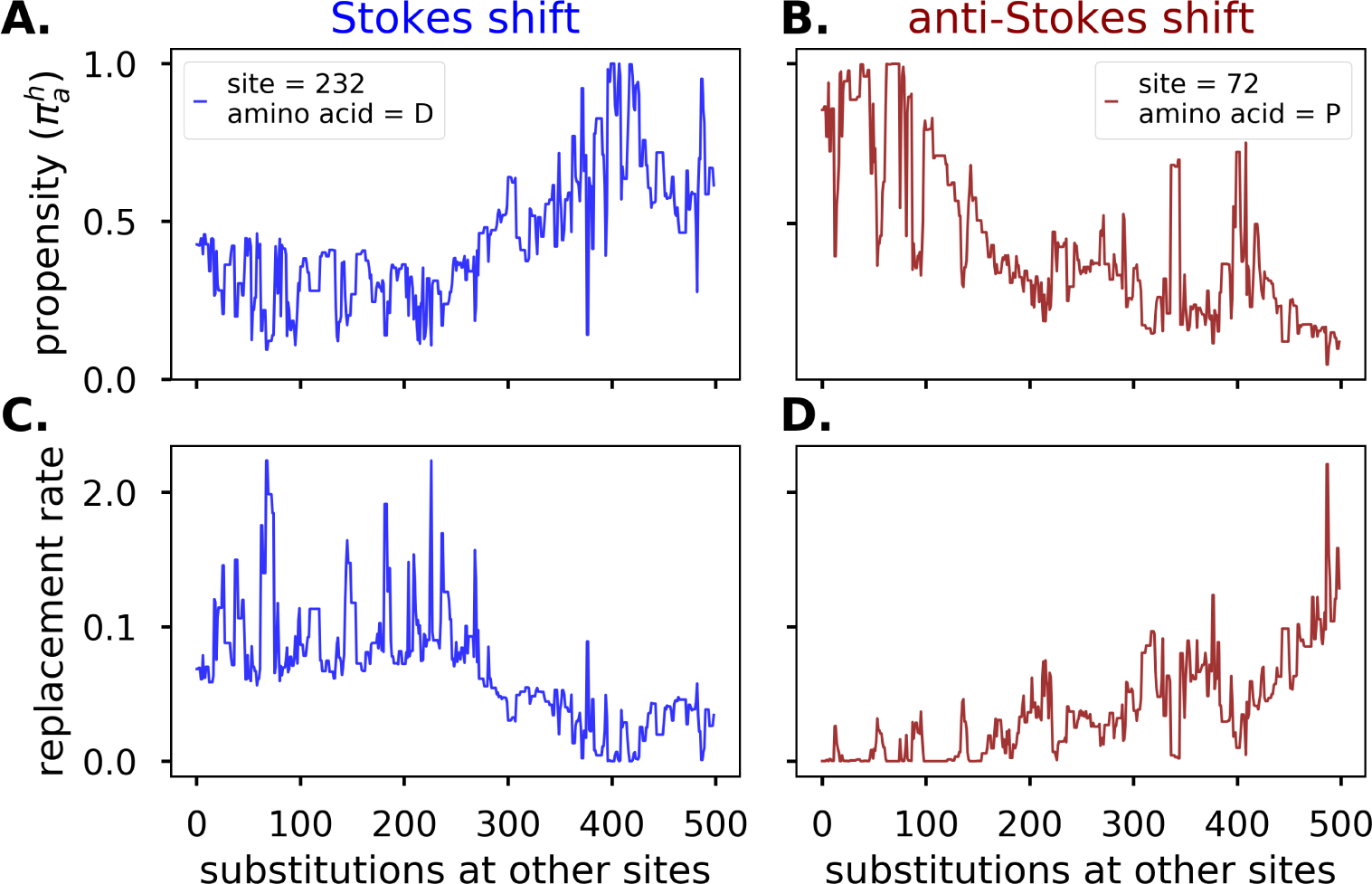
Neutral non-adaptive evolution can result in evolutionary Stokes and anti-Stokes shifts. Results are based on evolutionary simulation of the 1pek protein for 500 substitutions. Site 232 (A,C) undergoes an evolutionary Stokes shift whereas site 72 (B,D) undergoes an anti-Stokes shift. No substitutions occurred at either site, and the resident amino acids were aspartic acid (one letter code D) and proline (one letter code P) for sites 232 and 72, respectively. (A) and (B) plot the propensity of the resident amino acids as replacements occur at other positions in the protein. (C) and (D) show the expected rate of leaving the resident amino acid.

Changes in amino acid propensities are not directly observable in natural proteins. However, Popova *et al*. (2019) recently put forward that sustained increases (or decreases) in propensities may produce a detectable signal in natural protein alignments by causing a change in the rate of amino acid replacement. They suggest that the propensity of an amino acid is inversely related to its rate of replacement: if the propensity for the resident amino acid is high, then the rate of replacement should be low, and vice versa. Therefore, in addition to the amino acid propensities, we calculated the expected rate of replacement (the sum of the substitution rates to all single step neighbouring sequences that differ from the current sequence at the site of interest). Figure 1C and 1D confirms the predicted effect on the change in the rate of leaving the resident amino acids at sites 232 and For site 232, the increase in propensity with time (*i.e*., evolutionary Stokes shift) is accompanied by a decrease in the replacement rate (figure 1A&C). Similarly, the decrease in resident amino acid propensity at site 72 (*i.e*., evolutionary anti-Stokes shift), is accompanied by an increase in the rate of leaving (figure 1B&D). Importantly, these dynamics arose in a model where no adaptive changes are occurring. In other words, the proteins are evolving at mutation-drift-selection equilibrium with no change in protein structure or function. This shows that neither the evolutionary Stokes *nor* the anti-Stokes shift depends on any external change in selection on protein function or environment.

### Evolutionary anti-Stokes shifts are common under non-adaptive evolution

The previous result shows that the evolutionary anti-Stokes shift can occur without a change in protein function. But, as the anti-Stokes shift is a newly described phenomenon, it is unclear whether it is widespread or rare in natural proteins; Do evolutionary Stokes and anti-Stokes shifts occur with similar frequencies? To address this, we developed four metrics for quantifying these two phenomena. The metrics are described in detail in the Methods section and illustrated in figure 2. Briefly, the first metric (M1) is the slope of the linear model where *x* is time (measured in substitutions) and *y* is the propensity of the resident amino acid *a* at site 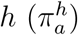. The slope is calculated over the amino acid residence time (*i.e*., from *i ≤ x ≤ j*, where *i* is the substitution when amino acid *a* first occupies the site and *j* is the last substitution). Metric two (M2) is the average change in the propensity of the resident amino acid over its residency time. As such, metrics M1 and M2 measure the *rate* at which propensities change over the time period where an amino acid is resident. In addition, we calculate a third metric (M3) - the difference in the average propensity of an amino acid while it is resident and the propensity of the same amino acid when it was first accepted-which estimates the *magnitude* of the change in propensity. Values of M1-3 greater than zero indicate an evolutionary Stokes shift, while values less than zero indicate an evolutionary anti-Stokes shift. Lastly, we define a more conservative metric, M4, which classifies an evolutionary Stokes shift only when M1-3 are all *>* 0, and an anti-Stokes shift when M1-3 are all *<* 0.

**Figure 2:**
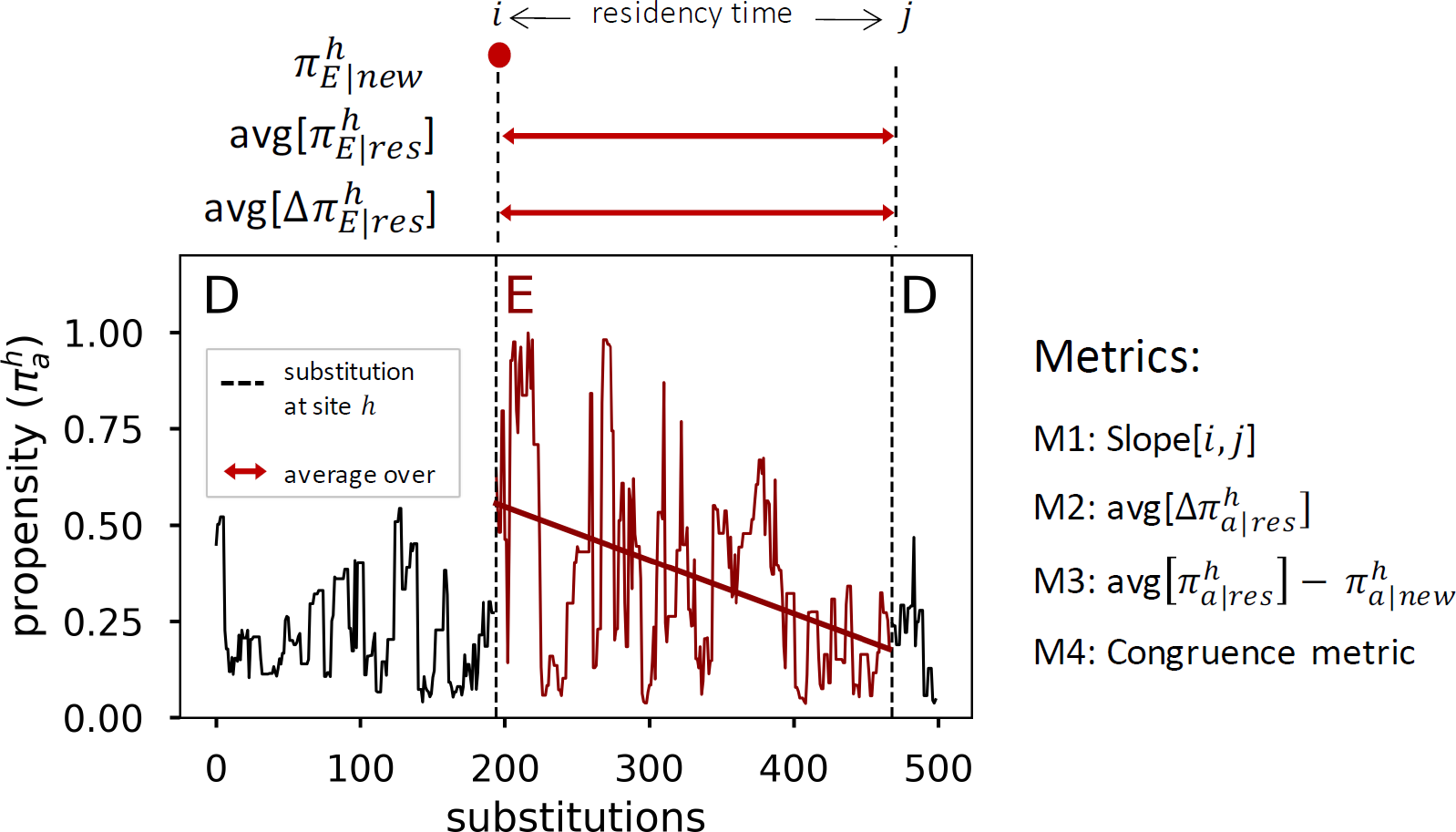
Description of metrics used to quantify evolutionary Stokes and anti-Stokes shifts. The example trajectory is based on site 82 of the 1pek protein. The site accepts two substitutions (vertical dotted lines) and the resident amino acid changes from D*→*E*→*D. For clarity, we focus on the dynamics following the acceptance of amino acid E. In general, metric 1 (M1) is the slope of the linear regression where *x* is the number of substitutions and *y* is the propensity of the resident amino acid *a* at site 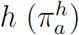 calculated over *i ≤ x ≤ j*; *i* is the substitution where amino acid *a* first occupies the site and *j* is the last substitution. Metric 2 (M2) is the average change in the propensity of the resident amino acid over its residency time (from *i* to *j*). Metric 3 (M3) is the difference between the average propensity of an amino acid while it is resident 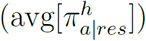and the propensity of the same amino acid when it was first accepted 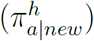. For M1-3, values *>* 0 indicate evolutionary Stokes shifts and values *<* 0 indicate evolutionary anti-Stokes shifts. Metric 4 (M4) is a congruence metric which classifies an evolutionary Stokes (or anti-Stokes) shift when all other metric values are *>* 0 (or *<* 0).

We found that an evolutionary anti-Stokes shift occurs following approximately half of all substitutions. The estimated proportion of substituted amino acids which experienced an evolutionary anti-Stokes shift (*P*_*anti−Stokes*_) were consistent across simulations under different protein structures and mutation models (Table S1). The consistency in *P*_*anti−Stokes*_ suggests that protein structures and mutation biases are not major determinants of evolutionary shifts in propensities. The proportions of *P*_*anti−Stokes*_ ranged from 0.46 to 0.57 across metrics M1-3 (Figure 3A). However, *P*_*anti−Stokes*_ estimated based on metric M4 were significantly less than 0.5 which is expected because of the more conservative requirements under M4. Furthermore, we found that, across all metrics M1-4, the proportion of substitutions that were followed by evolutionary anti-Stokes shifts (*P*_*anti−Stokes*_) were approximately equal to the proportion of substitutions that were followed by evolutionary Stokes shifts (*P*_*Stokes*_), suggesting that both phenomena occur with comparable frequencies (Figure 3B).

**Figure 3:**
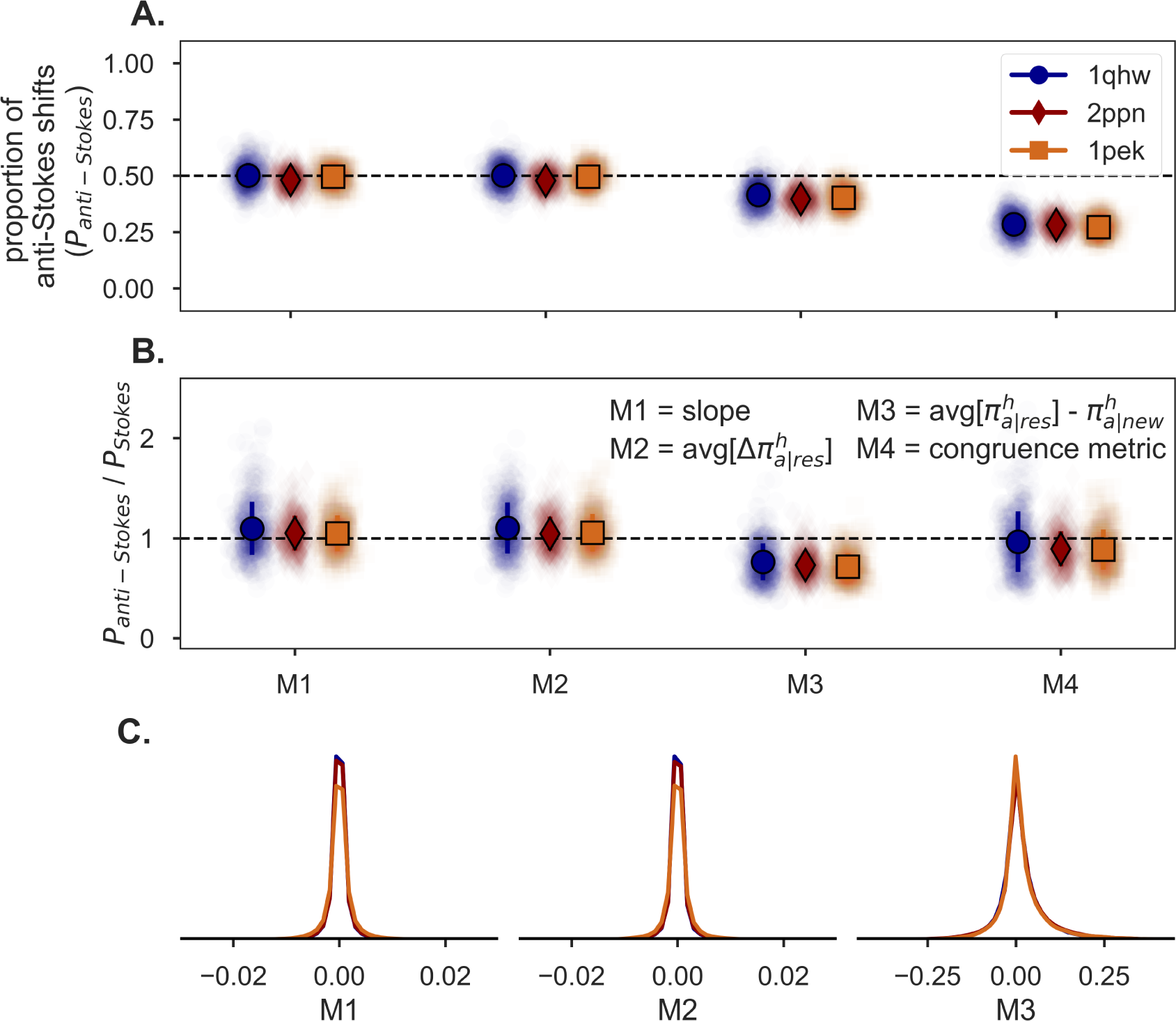
Evolutionary anti-Stokes shifts are common under non-adaptive evolution. (A) Approximately half of substitutions are followed by evolutionary anti-Stokes shifts based on metrics M1-3. The more conservative metric M4 estimates a slightly lower proportion (approx. 0.3). (B) Across all metrics M1-4, evolutionary Stokes and anti-Stokes shifts occur at similar frequencies (*P*_*anti−Stokes*_*/P*_*Stokes*_ *≈* 1). (C) The values of M1-3 are normally distributed and centered at zero which suggests that both the magnitude and frequencies of Stokes and anti-Stokes shifts are balanced.

Importantly, given the current formulations of the metrics, the estimates of both Stokes and anti-Stokes shifts could be comparable while the underlying dynamics may be considerably different. For example, the metrics would estimate similar values for the following scenarios: (1) a rapid increase (or decrease) in amino acid propensity followed by a longer period where the propensity of the amino acid remains high (or low), and (2) a more gradual increase (or decrease) in propensity over time. It may be the case that evolutionary Stokes shifts tend to occur soon after the acceptance of the substitution, while evolutionary anti-Stokes shifts may occur more gradually. To quantify whether propensity changes accelerated or decelerated over time, we compared the value of each metric M1-3 calculated over the first half of the amino acid residency (we label this as MX_1_) and the estimate over the second half (MX_2_), where X is either 1, 2, or 3 representing one of the three metrics. Specifically, we calculated (MX_2_ - MX_1_) / T_*res*_ where T_*res*_ is the amino acid residency time (measured in number of substitutions). We classified amino acids as undergoing an evolutionary Stokes or anti-Stokes shift based on the conservative metric M4. Using Welch’s t-test, we found significant differences in the average rates of change between Stokes and anti-Stokes shifts; however, the effect sizes were negligible (differences in means were *≤* 0.001, table S2).

Goldstein and Pollock (2017) observed that when a site experiences an evolutionary Stokes shift, not only does the propensity for the resident amino acid increase, but so does the propensity for physicochemically similar residues. For example, if V becomes newly resident at a site, then, over time, the propensity for similar amino acids (e.g. L) will likewise increase. This leads us to the question: does the decrease in propensity for the resident amino acid (an evolutionary anti-Stokes shift) imply a decrease in the propensities for similar amino acids? To address this, we grouped amino acids into bins of residues that tend to interchange rapidly and that tend to have similar chemical properties: [AST], [C], [DE], [FY], [GN], [HQ], [IV], [KR], [LM], [P], [W] (Susko and Roger, 2007). Rather than evaluating how the propensity for an individual resident amino acid changed over time, we tracked how the propensity of a group of amino acids changed (by summing the propensities of all amino acids within the group). We then applied metrics M1-4 to the summed group propensity. If evolutionary anti-Stokes shifts tend to affect individual amino acids, while Stokes shifts tend to occur for similar amino acids, then we would expect *P*_*anti−Stokes*_ ≠ *P*_*Stokes*_ based on the group analysis. However, we found that even when considering how the propensities for groups of similar amino acids changed *P*_*anti−Stokes*_ remained approximately equal to *P*_*Stokes*_ (figure S2). Overall, these results suggest that evolutionary anti-Stokes shifts are as common as Stokes shifts, and have similar dynamics.

### Evolutionary Stokes and anti-Stokes shifts occur at exposed and buried sites

It has long been observed that a site’s location in the protein influences its evolutionary dynamics. For globular proteins, surface residues are usually involved with protein function (*e.g*., binding affinity, enzymatic activity) with a preference for hydrophilic residues, while buried sites are usually occupied by hydrophobic residues and tend to evolve much slower (Yeh *et al*., 2013; Shahmoradi *et al*., 2014; Marcos and Echave, 2015; Echave *et al*., 2016). Popova *et al*. (2019) recently suggested that buried sites are more likely to undergo evolutionary Stokes shifts because of stability-constraints, while exposed sites are prone to senescence (decreases in amino acid favorability with time) due to changes in the environment external to the protein. We have shown that evolutionary anti-Stokes shifts can occur due to epistatic stability constraints without any external environmental or functional changes. Motivated by the results of Popova *et al*. (2019), we assessed whether some positions in the protein are more susceptible to evolutionary Stokes shifts, whereas others are more likely to undergo evolutionary anti-Stokes shifts. To do this, we examined the relationship between the metrics and two measures of the site’s location in the protein: relative solvent accessibility (RSA) and weighted contact number (WCN), both of which correlate significantly with substitutions rates in natural (Yeh *et al*., 2013; Shahmoradi *et al*., 2014; Marcos and Echave, 2015) and simulated proteins (Youssef *et al*., 2020). Specifically, exposed sites have higher substitution rates, higher RSA, and lower WCN as compared to buried sites. In line with these observations, we found a negative correlation between RSA and average residency time, and a positive correlation between WCN and average residency time (figure S3). However, we found no correlation between any of our metrics and RSA or WCN (figure S4 and S5). In this context (c.f. Popova *et al*. (2019)), the lack of correlations suggests there is no innate tendency for evolutionary Stokes or anti-Stokes shifts to be specifically associated with the location of sites in a protein.

While all sites are equally susceptible to undergoing evolutionary Stokes or anti-Stokes shifts, it remained unclear if the entailed dynamics were comparable. We were interested in assessing whether the site’s location in the protein might influence the rate of propensity changes. For example, if a deleterious substitution at a surface residue is easily compensated for (by adjustments at a small number of interacting sites), then we might expect a rapid increase in the resident amino acid propensity, and therefore a rapid evolutionary Stokes shift. Alternatively, if the site is highly connected, then the fixation of a deleterious amino acid at that site might require more adjustments at other positions, and, therefore, the Stokes shift might occur over a longer period of time. Across all metrics, we found no correlation between RSA and rate of propensity changes (figure S6) and, similarly, no correlation between WCN and rate of propensity changes (figure S7). This suggests that both exposed and buried sites undergo evolutionary Stokes and anti-Stokes shifts with comparable dynamics.

### Propensity shifts are consistent with random fluctuations

In the absence of selection, all mutations are neutral and are fixed (or lost) by the action of genetic drift, resulting in propensities that vary randomly over time (Wright, 1929). In contrast, our simulations take into account the action of selection with mutations conferring different fitness effects and, therefore, different fixation probabilities. Nonetheless, consistent with Goldstein (2011), we observed that approximately 90% of fixed nonsynonymous substitutions in our simulations were effectively neutral (i.e. where the selection coefficient from a current state *i* to a mutation *j* is |*s*_*ij*_| *<* 1*/*2*N*_*e*_; table S3). The high proportion of neutral fixations suggests that substitutions, and hence propensity fluctuations, are mainly driven by random genetic drift. Furthermore, the smooth symmetric M1-3 distributions centered at zero (figure 3C) are suggestive of random fluctuations in propensities. We were therefore interested in assessing if resident amino acid propensities tend to change randomly or whether they undergo phases of systematic increase and decrease over time. To address this, we conducted a mixed model analysis with a null model assuming random changes in propensities (see Methods). We did not find any evidence to reject the null random walk model. Interestingly, however, *all* p-values were *>* 0.95. This result is surprising since if propensities were varying randomly over time, then some sites should have rejected the null model just by chance. A potential explanation for the lack of fit (excess of high p-values), is if propensities changes were autocorrelated. Indeed, we observed a substantial negative autocorrelation in the differences in 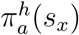 and 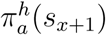 (table S4), implying that an increase in propensity tends to be followed by a decrease (and vice versa). This is perhaps not surprising since if the resident amino acid propensity decreases, then the site will either substitute away from the current amino acid or replacements will occur in other parts of the proteins increasing the propensity for that amino acid. Alternatively, as the propensity for a resident amino acid increases, there will be fewer ways for it to increase further than for it to decrease (for example, consider the dynamics when propensity is equal to one).

Lastly, we were interested in assessing whether random fluctuations in propensities could result in *P*_*Stokes*_ and *P*_*anti−Stokes*_ comparable to those observed in our simulations. To do this, we simulated 500 bounded random walks (between 0 and 1) of amino acid propensities with step sizes drawn from a normal distribution with mean (*µ*=0) and standard deviation (*σ*= 0.1) estimated from the step sizes observed in the stability-constrained simulations (figure S8). Then, we applied metrics M1-4 to the random walk simulations in order to estimate the proportion of evolutionary Stokes and anti-Stokes expected when propensities vary randomly. We found that the proportion of evolutionary Stokes and anti-Stokes shifts from stability-constrained simulations were consistent with the proportions estimated under a bounded random walk model (figure S9). Overall, these results suggest that the dynamics of propensities are likely not too different from a random walk at equilibrium, and that random fluctuations in resident amino acid propensities will occasionally lead to increases in propensities over time (evolutionary Stokes shifts) and decreases in propensities over time (evolutionary anti-Stokes shifts).

### Most stabilizing substitutions are permissive and most destabilizing substitutions are restrictive

We have shown that the evolutionary Stokes and anti-Stokes shifts are both common within a system evolving under selection for stability with no adaptive changes. We next turn to the underlying mechanisms that give rise to these dynamics. Important questions about how substitutions impact resident amino acids propensities remain unanswered: Do substitutions tend to favourably impact some sites (by increasing their resident amino acid propensities) while simultaneously disadvantaging other sites (by decreasing their resident amino acid propensities)? Or does a substitution tend to impact the propensity of resident amino acids similarly across the protein? If so, what explains the observed balance between *P*_*Stokes*_ and *P*_*anti−Stokes*_?

A consequence of stability-mediated epistasis is that any change in the protein sequence will cause shifts in resident amino acid propensities at other sites. For example, when a substitution occurs, so that the sequence changes from *s*_*x*_ *→ s*_*x*+1_, the fitness and propensity landscapes at most other sites in the protein will subsequently change. In figure 4A, the grey dots represent the change in the propensity of the resident amino acid at each site following a substitution 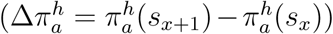. The red dots represent the change in the propensity of the resident amino acid at the substitution site, and therefore a change in the resident amino acid from 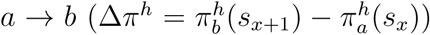. We found that the effect of a substitution on resident amino acid propensities is usually unbalanced. In other words, substitutions either favourably impact most sites (so that the proportion of sites with negative 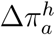 is less than 0.5), or decrease the propensity of the resident amino acid at most sites (so that the proportion of sites for with negative 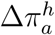 is greater than 0.5). Surprisingly, stabilizing substitutions (ΔΔ*G <* 0) were associated with decreases in propensities of resident amino acids at most sites, while destabilizing substitutions (ΔΔ*G >* 0) rendered the resident amino acids at most sites more favorable (figure 4B).

**Figure 4:**
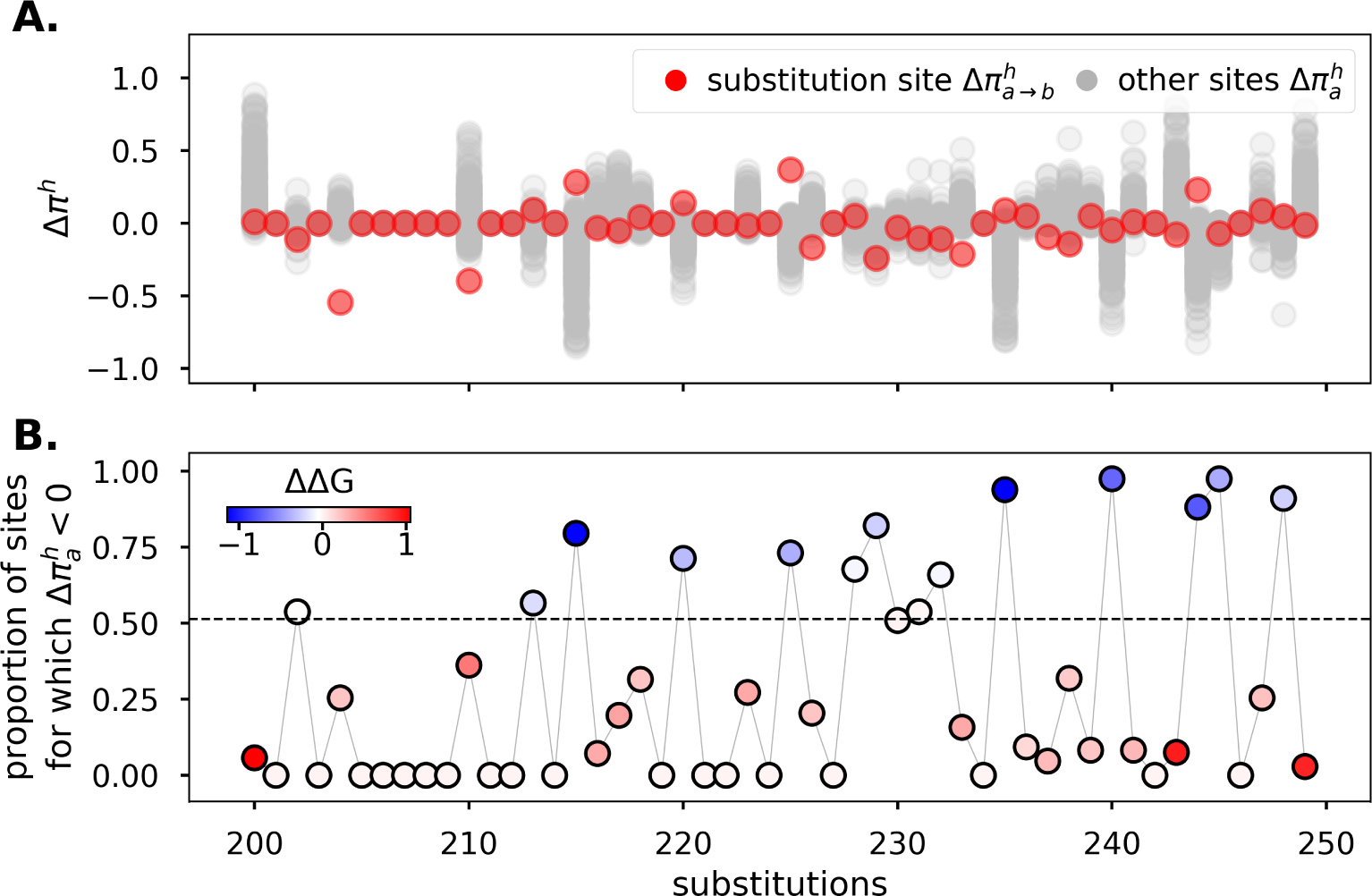
Stability-mediated epistasis between sites results in changes in resident amino acid propensities as substitutions occur in the protein. (A) Following an amino acid replacement at one position in the protein, so that the sequence changes from *s*_*x*_ *→ s*_+1_, the propensity of the resident amino acids at all sites will subsequently change. The grey dots are the changes in the propensities of the resident amino acids at each site following a substitution, 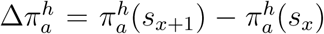. The red dots are the change in the propensity of the resident amino acid at the substitution site, and therefore a change in the amino acid from 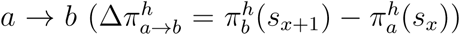. (B) Stabilizing substitutions (ΔΔ*G <* 0) result in higher proportions of 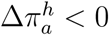. In other words, they result in a decrease in the propensity of amino acids at most other sites. In contrast, destabilizing substitutions (ΔΔ*G >* 0) result in lower proportions of 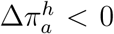. Dotted line is the average proportion of 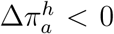 over all substitutions (average = 0.51).

This result initially appears counter-intuitive. To clarify, let us consider examples of the dynamics following the fixation of a stabilizing and a destabilizing substitution observed within our simulations (figure 5). First, consider the dynamics following a stabilizing substitution. Let *s*_1_ be the initial protein sequence with fitness = 0.999 and stability (-Δ*G* = 4.041; figure 5A). A substitution occurred so that the sequence changed from *s*_1_ *→ s*_2_, that increased the overall stability of the protein with a minor improvement in fitness. As a result of this substitution, the fitness landscapes at most sites changed. We focus on the fitness landscape at site 145 before (*s*_1_) and after (*s*_2_) the change in the background sequence. Given that sequence *s*_2_ is more stable, a destabilizing mutation at site 145 has relatively lower impact on the fraction of correctly folded proteins at equilibrium (i.e., fitness). Thus, the effect of the background sequence having higher stability is that the fitness landscape at site 145 becomes more uniform (figure 5B). How does the change in the fitness landscape induce a change in the propensity of the resident amino acid? Since a larger number of amino acids can now occupy the site with little or no detriment to protein fitness, the propensity landscape will similarly become more uniform (figure 5C). Amino acids like R, N, and P that had low propensity in the context of background sequence *s*_1_, are more likely to be observed given the “stability-buffered” sequence *s*_2_ (figure 5C). Propensities are the expected equilibrium frequencies of each amino acid given the current background sequence; they must, therefore, sum to one. The increase in the propensity of some amino acids (*e.g*., R, N, and P) will cause a decrease in the propensity of the resident amino acid (K in this example; figure 5). The opposite relationship was observed following the fixation of a destabilizing mutation (figure 5D): The fitness and propensity landscapes became less uniform (figure 5E&F), with fewer amino acids having non-zero propensities. This resulted in an increase in the propensity for the resident amino acid following the destabilizing substitution (figure 5F).

**Figure 5:**
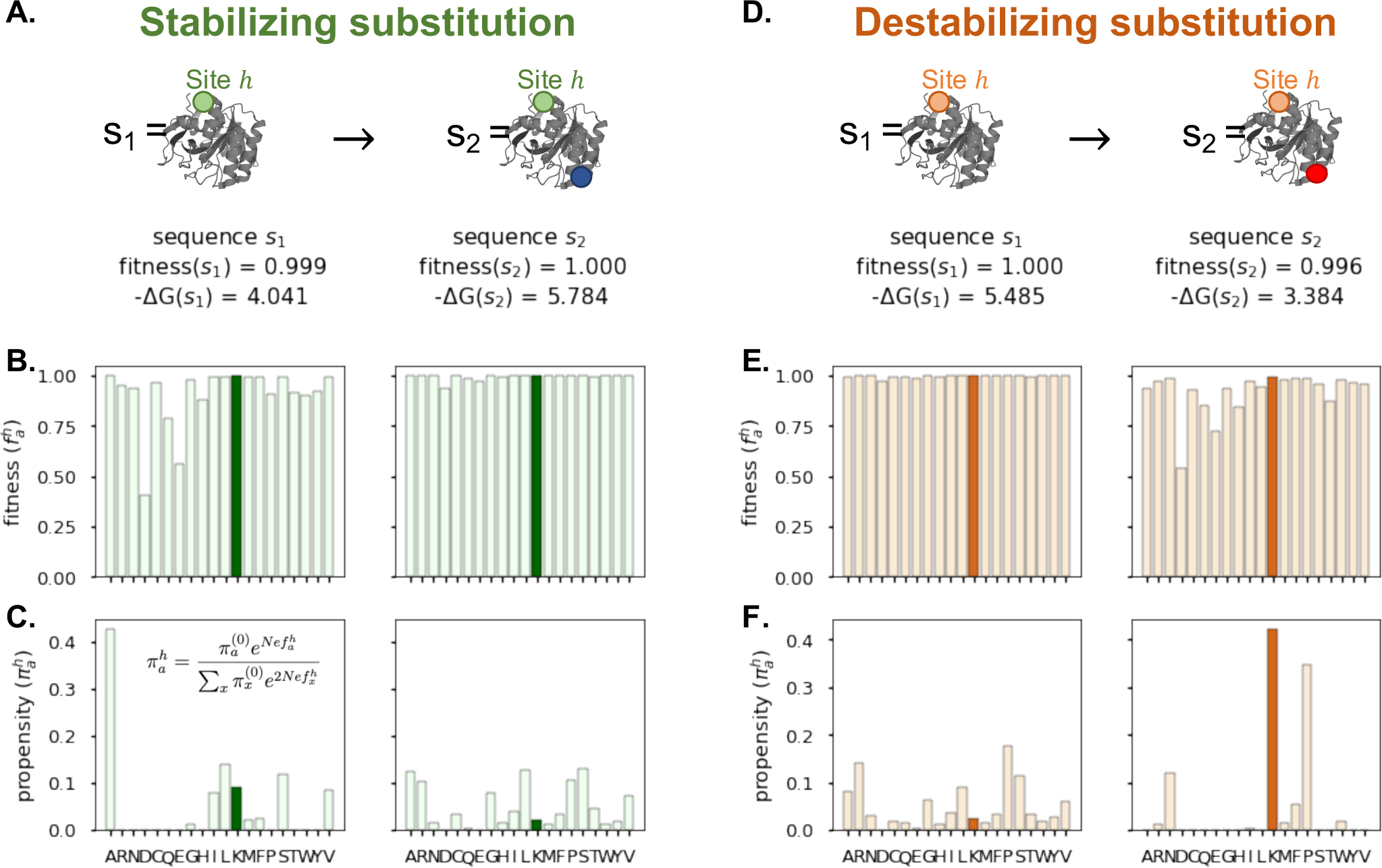
Epistatic dynamics following the fixations of stabilizing (A,B,C) and destabilizing (D,E,F) substitutions. (A) Let *s*_1_ be the initial protein sequence, and *s*_2_ be the sequence following the acceptance of a stabilizing substitution (blue dot). Given the “stability-buffered” background sequence *s*_2_, deleterious mutations which would not have been fixed in the context of background sequence *s*_1_ are now more likely to be fixed (e.g. R, N,P). (B) The fitness landscape at a non-substituted site *h* becomes more uniform because of the increase in overall protein stability. (C) Similarly, the propensity landscape becomes more uniform. The fitness and propensity of the resident amino acid is shown in dark green. (D, E, F) are the respective plots following the fixation of a destabilizing substitution (red dot). The fitness and propensity landscapes at the non-substituted site become less uniform.

To quantify the uniformity of a landscape, we calculated its Shannon entropy H^*h*^(*s*) (see Methods section for detail). Entropy is maximized when the landscape is uniform (*i.e*., all amino acids have equal frequencies), and is at a minimum (= 0) when only one amino acid is observed at the site. Note that entropy of fitness and propensity landscapes are highly correlated (figure S10). The fitness landscape describes fitnesses of nearby sequences, whereas the propensity landscape considers how frequently nearby sequences are explored via mutation and substitution. We, therefore, report the entropy of the propensity landscapes, although similar results are expected based on the fitness landscapes. We found that at higher stability values (lower Δ*G*) the propensity landscapes at most sites tend to be more uniform compared to at lower stability values (figure 6A). This suggests that as the protein becomes more stable, most amino acids tend to have similar impacts on fitness and are, therefore, equally likely to be observed at a site. Next, we were interested in assessing the impact of a substitution on the uniformity of the landscapes at other sites in the protein. To do this, we calculated the change in landscape entropy following the acceptance of a substitution, ΔH^*h*^ = H^*h*^(*s*_*x*+1_) - H^*h*^(*s*_*x*_). A change from a more uniform to a more rugged landscape (where a smaller number of amino acids have non-zero propensities) will result in a negative ΔH^*h*^. In contrast, a positive ΔH^*h*^ indicates a change towards a more uniform landscape. We considered a substitution as permissive if it causes a flattening in the propensity landscape at most sites (*i.e*., a positive average ΔH). A restrictive substitution is one where following its acceptance, the landscapes at most sites permit fewer amino acids (*i.e*., a negative average ΔH). We found a strong positive correlation between the stability effect of a substitution (ΔΔ*G*) and its influence on the fitness landscapes at other sites (figure 6B). Figure 6C reports the average proportions of the different types of substitutions estimated based on three protein-specific simulations. Consistent with the results reported in figure 5, stabilizing substitutions - by increasing the overall stability of the protein - provide a “stability-buffered” background so that slightly destabilizing substitutions are more likely to be fixed, expanding the space of potential evolutionary paths. In contrast, destabilizing substitutions restrict potential evolutionary paths. These results are in line with experimental work by Gong *et al*. (2013), which found that stabilizing substitutions permit otherwise inaccessible mutations.

**Figure 6:**
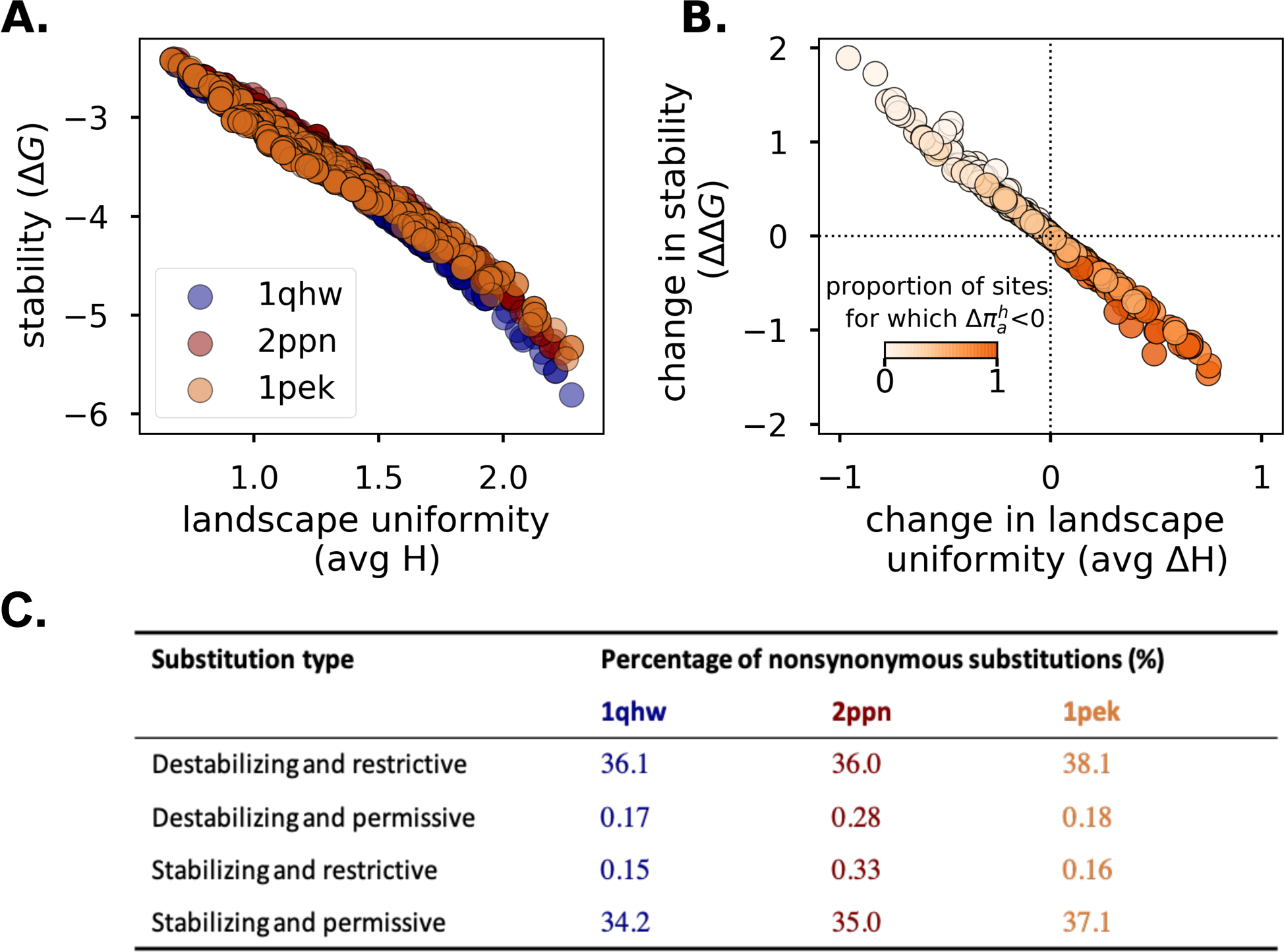
Most stabilizing substitutions are permissive and most destabilizing substitutions are restrictive. (A) The relationship between protein stability (Δ*G*) and landscape uniformity, measured as the entropy of the propensity landscape averaged over all sites in the protein (avg H). (B) The relationship between the stability effect of a substitutions (ΔΔ*G*) and the resulting average change in landscape uniformity (avg ΔH). Color bar represents the proportion of sites for which the propensity for the resident amino acid decreased 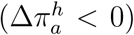. Positive avg ΔH values imply that, on average, the landscapes became more uniform. Therefore, the substitution is deemed permissive. Negative avg ΔH are indicative of restrictive substitutions. Plotted results are based on a single simulation of the 1pek protein. (C) The percentages of different types of substitutions for each of three proteins (1qhw, 2ppn, and 1pek). Percentages are calculated from 500 protein-specific trials

These results are also consistent with previous theoretical predictions by Cherry (1998), and Goldstein (2011). Cherry (1998) observed that, given a saturating fitness function, the distribution of potential mutational fitness effects (*s*_*ij*_ = *f*_*j*_ *− f*_*i*_ where *i* and *j* are the wildtype and mutant alleles respectively) will be related to the current fitness of the organism (or protein): the distribution of fitness effects broadens as fitness decreases. In the model used here, fitness (the probability of folding) is a saturating function of protein stability (Δ*G*, equation 4). We find that at higher stability values (lower Δ*G*) the propensity landscapes tend to be more uniform (figure 6A). This suggests that the magnitude of the fitness effects for potential mutations are smaller at higher stability values compared to the distribution when the background sequence is less stable. Second is the prediction that at equilibrium the proportions of deleterious substitutions will be balanced by the proportion of beneficial substitutions. We found that the proportion of stabilizing (beneficial) substitutions balanced those that were destabilizing (deleterious) in our simulations (figure 6B). Taken together, these predictions shed light onto the observed balance between *P*_*Stokes*_ and *P*_*anti−Stokes*_. At mutation-drift-selection equilibrium we expect that *P*_*stabilizing*_ = *P*_*destabilizing*_ (figure 6B; Cherry (1998); Goldstein (2011)). We have shown that stabilizing substitutions result in a decrease in the propensity of the resident amino acids at most sites (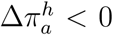, because of a flattening in the site-specific fitness landscapes), while destabilizing substitutions increase resident amino acids propensities at most sites (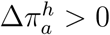, figure 5 & 6A). This suggests that the proportion of 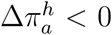 should be equal to the proportion of 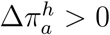 (figure S8). Since evolutionary Stokes and anti-Stokes shifts are the result of such changes in propensities, we would expect that at equilibrium *P*_*Stokes*_ *≈ P*_*anti−Stokes*_.

## Discussion

We have examined evolutionary dynamics under a nonadaptive stability-constrained model of protein evolution. Consistent with previous observations, we found that as proteins become more stable (lower Δ*G* values), the fitness effects of most mutations diminished compared to the fitness effects of the same mutation at lower stability (higher Δ*G* values). This suggests that substitutions which increase stability will make more mutations accessible to the protein, thereby expanding the space of potential evolutionary paths. In contrast, destabilizing substitutions will tend to limit potential evolutionary trajectories.

Previous studies observed that functionally important residues are often destabilizing (Schreiber *et al*., 1994; Nagatani *et al*., 2007; Wang *et al*., 2002; DePristo *et al*., 2005), suggesting a trade-off between function and stability. This leads to the hypothesis that highly stable proteins are more readily adaptable to new functions compared to less stable proteins since they are more likely to accept destabilizing yet functionally beneficial substitutions. We suggest that more stable proteins, all other things being equal, may be more adaptable, not only because they can tolerate destabilizing yet functionally beneficial substitutions, but also because they are more apt to explore neighboring regions of sequence space. At low protein stability fewer mutational paths are permissible (since most changes are deleterious), resulting in longer waiting times between substitutions and hence fewer opportunities to explore sequence space and adapt to other functions. However, it is important to note that selection on other properties of proteins, such as their expression level and the cost of translation error (Drummond *et al*., 2005), can also influence their rate of evolution. Therefore, the relationship between evolvability and stability of proteins in nature is likely to reflect the complex interplay of multiple factors.

As more (or fewer) mutations become accessible, the propensity (i.e., the equilibrium frequencies given a particular background sequence) for the currently resident amino acid at a site will consequently change. By expanding accessible paths, stabilizing substitutions tended to result in a decrease in the propensity for the resident amino acids at most sites, while destabilizing substitutions frequently increased propensities (figure 4B & 6A). Shifts in resident amino acid propensities were occasionally consistent with an evolutionary Stokes shifts (Pollock *et al*., 2012), where the propensity for an amino acid increases over time due to compensatory adjustments at other sites in the protein. In other instances, we observed the opposite trend where the propensity of a resident amino decreased with time, an evolutionary anti-Stokes shift. In our simulations, both evolutionary Stokes and anti-Stokes shifts were caused entirely by stability-mediated epistasis and not by any external changes in protein function or environment. Alternatively, Popova *et al*. (2019) recently observed that the fitness of the resident amino acid at a site may decrease with time since substitution. They attributed the decrease in amino acid fitnesses to external changes in the protein’s environment (*e.g*., host immune response) and termed this trend senescence. This highlights the main difference between the notion of an evolutionary anti-Stokes shift and the concept of senescence: evolutionary anti-Stokes shifts are a result of non-adaptive processes mediated by the emergent property of marginal stability, whereas senescence is a consequence of an adaptive response to some change in the protein’s external environment.

Alternatively, propensity shifts may be viewed as dynamics that arise due to the protein adapting to internal, rather than external, changes. In this sense, neighbouring sites may “compensate for” or “adapt to” a deleterious substitution that occurred at an interacting site. However, our results challenge even this more localized interpretation of adaptation. Evolutionary dynamics at equilibrium are predominantly driven by random drift where the vast majority of substitutions (approximately 90%, table S3) are nearly neutral with |*s*_*ij*_| *<* 1*/*2*N*_*e*_. If drift were the only evolutionary force acting on a gene (e.g. pseudogene), then allele propensities are expected to vary randomly over time. We found that propensity changes are largely consistent with random fluctuations in propensities that may occasionally favour or disfavour a resident amino acid at a site. Furthermore, if evolutionary Stokes shifts are the result of local co-adaptation, then we would expect a higher fraction of substitutions at neighbouring sites compared to the proportion when an anti-Stokes shift occurs. However, we found no evidence in support of this; the fraction of substitutions at neighbouring sites were comparable for both evolutionary Stokes and anti-Stokes shifts (figure S11).

We see that a major advantage of the thermodynamic stability model used here, and previously (Goldstein, 2011; Pollock *et al*., 2012; Goldstein and Pollock, 2017), is that it provides a plausible nonadaptive null model for protein evolution. Such stability-informed models of protein evolution have previously been used to discredit adaptationist claims about the necessary trade-offs between protein function and stability (Taverna and Goldstein, 2002; Goldstein, 2011), and protein function and foldability (Govindarajan and Goldstein, 1996). “Despite the seduction of adaptive rationalizations”, to quote one of the original authors of this model, “neutral evolutionary dynamics remains the null model that must first be rejected” (Goldstein, 2011). Our observation that amino acid propensities may decrease over time in the absence of external environmental changes does not preclude that environmental shifts could render resident amino acid less favourable. Rather our simulations demonstrate that decreases in propensities can occur in the absence of environmental changes, and therefore that their mere occurrence should not, on their own, be taken as conclusive evidence of adaptations to external environmental changes.

## Methods

### Protein description

We simulated the evolution of three proteins with PDB codes 1qhw, 2ppn, and 1pek. These protein structures are described in detail in Youssef *et al*. (2020). The proteins differ in structure, function, length, and contact density. The 1qhw protein is a phosphatase, the 1pek protein is a proteinase, and the 2ppn protein is an isomerase. The 1qhw protein is 300 amino acid residues long, 1pek is made of 297 amino acids, and the 2ppn protein comprises 107 residues. The 1pek protein was the most densely packed with an average number of contacts per site of 8.4 compared to 7.5 for the 1qhw protein and 6.9 for the 2ppn protein. During the simulations, we used the nucleotide frequencies (*π*_*n*_) and transition/transversion rate (*κ*) parameters estimated from multiple sequence alignments for the corresponding protein used in Youssef *et al*. (2020). The mutation parameters (*κ, π*_*A*_, *π*_*C*_, *π*_*G*_, *π*_*T*_) were set equal to (4.37, 0.21, 0.32, 0.28, 0.20) for the 1qhw protein; (0.90, 0.19, 0.35, 0.56, 0.21) for the 1pek protein; and (2.50, 0.27, 0.24, 0.29, 0.19) for the 2ppn protein.

### Evolutionary model

The evolutionary process is based on the mutation-selection (MutSel) framework (Halpern and Bruno, 1998). We assume a Wright-Fisher population with fixed effective population size (*N*_*e*_) evolving under a weak mutation, strong selection regime so that only a single variant exists in the population at any time point. The probability of a particular mutation *b* going to fixation in a diploid population currently fixed at variant *a* is calculated as

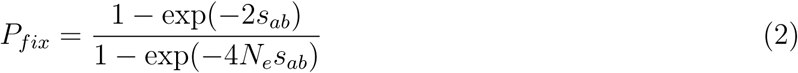

where *s*_*ab*_ = *f*_*b*_ *−f*_*a*_ is the relative fitness effect of mutant *b* (Kimura, 1962). We model the substitution process as a continuous-time Markov chain which is specified by the instantaneous rate matrix *Q* with elements

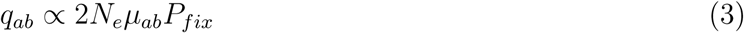

where *q*_*ab*_ is the substitution rate from *a* to *b* which depends on the mutation rate (*µ*_*ab*_) and the fixation probability (*P*_*fix*_). Mutations arise at the DNA-level following the HKY model (Hasegawa *et al*., 1985) allowing only single nucleotide changes. Selection is assumed to act on the final protein product, and therefore all synonymous codons have the same fitness. We assumed a fixed *N*_*e*_ = 100.

We initiated each simulation at a randomly generated amino acid sequence. Then, we used the algorithm outlined in table S5 to obtain protein sequences with fitness values *≥* 0.99 given the corresponding structure. Following this equilibration phase, we evolve the equilibrated sequence for 500 substitutions while keeping track of the site-specific fitness landscapes at all sites. The reported results are based on the post-equilibration phase. We generated 500 protein-specific replicates.

### Stability model

We use the same stability model outlined in Goldstein (2011), Pollock *et al*. (2012), Goldstein and Pollock (2017), and Youssef *et al*. (2020). For a detailed description of the model derivation see section “The protein model” in Goldstein (2011). Briefly, we assume that the fitness is equal to the probability of an amino acid sequence being in the native (folded) structure at thermodynamic equilibrium, which is a function of the stability (Δ*G*) of the sequence.

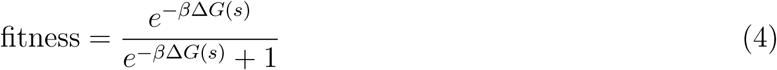

where 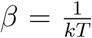, *k* is the Boltzmann constant, *T* is the absolute temperature, and Δ*G*(*s*) is stability of sequence *s* measured as the difference in free energy between the folded and the unfolded states. The free energy of a sequence in a given structure is approximated as the sum of pairwise potentials (from Miyazawa and Jernigan (1985)) for amino acids in contact. Residues are considered to be in contact if the C_*β*_ atoms are within 7°Å of each other. If the amino acid present is glycine, distance is considered with reference to the C_*α*_ atom.

### Amino acid propensities

Suppose that for a simulation trial we observed *s*_1_ *→ s*_2_ *→* … *→ s*_500_ where the *s*_*x*_’s are the codon sequences realized during the simulations, and *s*_*x*_ and *s*_*x*+1_ differ by a single nucleotide substitution (synonymous or nonsynonymous). Given each *s*_*x*_, we can calculate the fitness landscape at site *h*, given the rest of the sequence is held constant, as 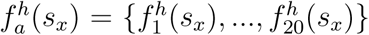. We use the fitness values to calculate the amino acid stationary frequencies using equation (1). We calculate 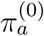 as the sum over the neutral stationary frequencies for synonymous codons for each amino acid. The neutral frequency for a codon made up of nucleotide triplet *ijk* will be proportional to *π*_*i*_*π*_*j*_*π*_*k*_.

### Description of metrics used to quantify evolutionary Stokes and anti-Stokes shifts

We define four metrics to quantify shifts in propensities. First, let the residence time of an amino acid (T_*res*_) be the time period between *i* and *j*, where *i* is the substitution when amino acid *a* first occupies the site and *j* is the last substitution. The first metric (M1) is the slope of the linear model over T_*res*_ where *x* is time (measured in substitutions) and *y* is the propensity of the resident amino acid *a* at site 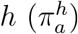.

Metric two (M2) is the average change in the propensity of the resident amino acid over its residency time. Following each substitution we calculate the change in propensity as

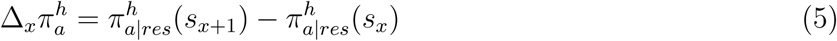

where 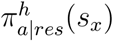 is the propensity of the resident amino acid given the background sequence present at substitution *x*. M2 is then the average calculated as

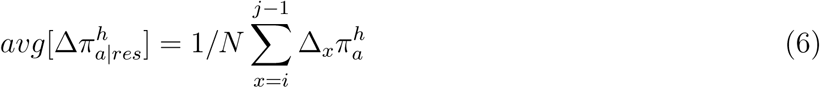

where *N* = *j − i*.

Pollock *et al*. (2012) observed that following an evolutionary Stokes shift “the inherent propensity for [an] amino acid at that position will be, on average, higher than it was when the substitution occurred”. We, therefore, define a third metric, which is perhaps the most consistent with the definition provided by Pollock *et al*. (2012). Metric three (M3) is the difference in the average propensity of an amino acid while it is resident 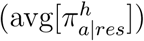 and the propensity of the same amino acid when it was first accepted 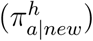.

For M1, M2, and M3, values greater than 0 are suggestive of an evolutionary Stokes shift and values less than 0 are indicative of evolutionary anti-Stokes shifts. Lastly, we define a more conservative metric, M4, which classifies an evolutionary Stokes shift only when all three metrics indicate a Stokes shift (all M1-3 values are *>*0), and an anti-Stokes shift when M1-3 are all *<* 0. Figure 2 provides a visual representation of the metrics.

### Quantifying the uniformity of a landscape

We use the entropy of a propensity landscape as a measure of its uniformity. We calculate entropy as

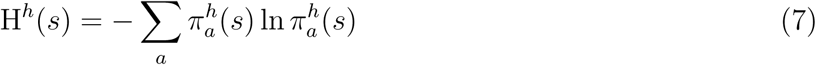

where 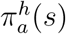 is the propensity of amino acid *a* at site *h* given background sequence *s*. The entropy is maximized when all amino acids are equally likely, and is minimized (= 0) when only a single amino acid is observed. To determine how the landscapes change in response to changes in the background protein sequence, we compared the entropy before and after the substitution

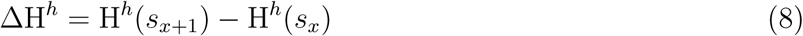

We classified a substitution as permissive if the average ΔH across all sites was positive, and restrictive if the average ΔH was negative.

For all results described in this study, we only considered the dynamics when a residue was accepted and subsequently replaced within the time-frame of the simulation. However, we repeated the analyses with the inclusion of partial windows (where for example an amino acid is accepted during the simulation but the simulation ends prior to its replacement) which revealed similar results with respect to the proportion of evolutionary Stokes and anti-Stokes shifts (figure S12).

### The rate of amino acid replacement

Popova *et al*. (2019) recently observed that changes in amino acid propensities are accompanied by changes in the relative rates of leaving the resident amino acid. Amino acids that have high fitness values, are more likely to occupy the site (have high 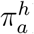), and will have a low rate of being replaced. Conversely, sites with low fitness benefit are less likely to be present at the site (low 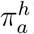), and will have a high rate of being replaced. Therefore, in addition to amino acid propensities, we looked at the replacement rates over time. We calculate the rate of leaving the resident amino acid at a site *h* as the sum of the transition rates (using equation (3)) over all sequences that differ from the current sequence by a single nucleotide and have a different amino acid at site *h*.

### Mixed linear model analysis

In order to assess if amino acid propensities shifts were consistent with random fluctuations we fitted the data to a mixed linear model of the form

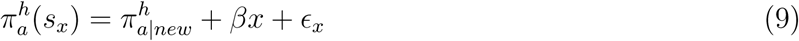

where *ϵ*_*x*_ *∼* N(0,*σ*^2^) and 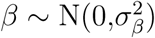. We tested a null model assuming random shifts in ropensities where 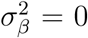 against an alternative model where 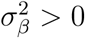.

### Code availability

All code used to simulate, analyze, and plot data has been uploaded and is freely available from https://github.com/noory3/antiStokes shifts.

