## Supplemental Material for "The ups and downs of amino acid co-evolution: evolutionary Stokes and anti-Stokes shifts"

**Supplementary material:**

**Table S1.** Proportion of substituted amino acids which experienced an evolutionary anti-Stokes shift calculated based on four metrics (M1-4). The metrics are described in the Methods section and illustrated in Figure 2, in the main document. Reported are the average proportions based on 500 protein-specific simulations for three protein structures (1qhw, 1pek, and 2ppn).

| | 1qhw | 1pek | 2ppn |
|---|---|---|---|
| M1<br>Slope[i, j] | 0.47 | 0.48 | 0.46 |
| M2<br>avg[ $\Delta\pi^{h}{}_{\{a\|res\}}$ ] | 0.47 | 0.48 | 0.46 |
| M3<br>avg[ $\Delta\pi^{h}{}_{\{a\|res\}}$ ] - $\Delta\pi^{h}{}_{\{a\|new\}}$ | 0.56 | 0.57 | 0.55 |
| M4<br>congruence metric | 0.31 | 0.31 | 0.32 |

**Table S2**. Differences in rate of acceleration (or deceleration) of evolutionary Stokes and anti-Stokes shifts. Reported are the P-values based on Welch's t-test. Substitutions are classified as undergoing an evolutionary Stokes or anti-Stokes shift based on metric M4. $T_{res}$ is the amino acid residency time measured as the number of substitutions during which an amino acid was resident at the site. $MX_1$ represents the value of each metric (X = 1,2, or 3) calculated over the first half of the amino acid residency, and $MX_2$ is the value of the same metric calculated over the second half.

| | 1qhw | | 1pek | | 2ppn | |
|---|---|---|---|---|---|---|
| | Difference in means | P-value | Difference in means | P-value | Difference in means | P-Value |
| $M1_2$ - $M1_1$ / $T_{res}$ | -6.5e-5 | < 0.001 | -7.4e-5 | < 0.001 | -8.2e-5 | < 0.001 |
| $M2_2$ - $M2_1$ / $T_{res}$ | 0.001 | < 0.001 | 0.001 | < 0.001 | -8.4e-5 | < 0.001 |
| $M3_2$ - $M3_1$ / $T_{res}$ | -6.7e-5 | < 0.001 | -7.8e-5 | < 0.001 | -8.4e-5 | < 0.001 |

**Table S3.** Proportion of nonsynonymous (and all) substitutions which were deleterious, neutral, or beneficial from simulations of three proteins with PDB structures: 1qhw, 2ppn, 1pek.

| | Deleterious $s_{ij} < -1/2N_e$ | Neutral $-1/2N_e < s_{ij} < 1/2N_e$ | Beneficial $s_{ij} > 1/2N_e$ |
|---|---|---|---|
| 1qhw | 0.05 (0.02) | 0.89 (0.94) | 0.06 (0.03) |
| 2ppn | 0.06 (0.03) | 0.88 (0.93) | 0.06 (0.03) |
| 1pek | 0.05 (0.03) | 0.90 (0.93) | 0.05 (0.03) |

**Table S4**. Autocorrelation of the differences in propensities, $\pi_a^h(s_x)$ - $\pi_a^h(s_{x-1})$, for $i \leq x \leq j$ where $i$ is the substitution when amino acid a first occupies the site and $j$ is the last substitution. Reported are the ranges of the average lag 1 autocorrelation over all sites.

| Protein | min | max |
|---|---|---|
| 1qhw | -0.13 | -0.08 |
| 2ppn | -0.12 | -0.09 |
| 1pek | -0.16 | -0.12 |

**Table S5.** Algorithm for obtaining sequences with high fitness (> 0.99) values**,** from Youssef et al 2020.

1. Start at random amino acid sequence
2. Calculate the site-specific fitness landscape at all sites
3. If a single step uphill move is possible (i.e. beneficial mutation), then randomly choose the next substitution from the set of single amino acid changes that will increase fitness
4. If no uphill move is possible (i.e. local maximum), then randomly choose 20 sites and substitute them to the fittest amino acid at that site
5. Repeat 2-4 until fitness is greater than 0.99

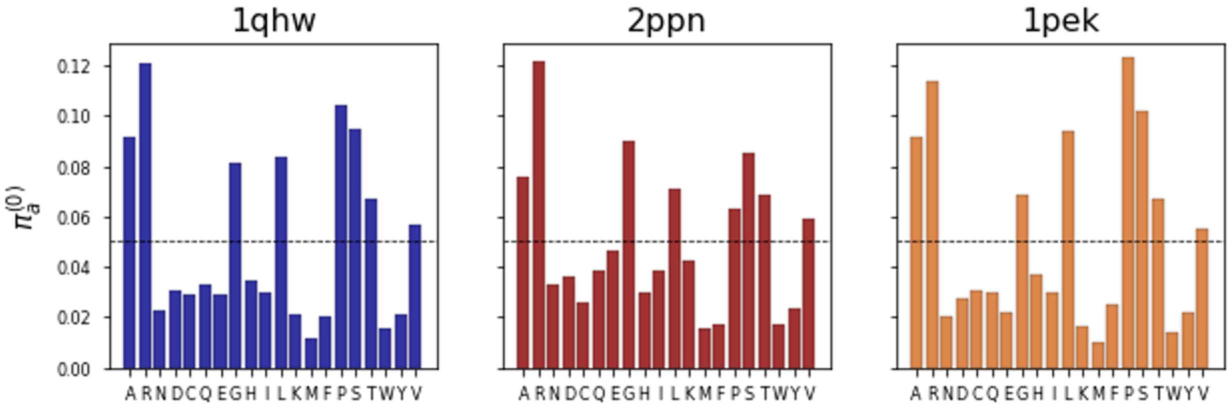


**Figure S1.** The expected amino acid frequencies in the absence of selection but accounting for underlying mutational biases. The dotted line represents the expected frequency values in the absence of mutational biases and assuming all amino acid have the same number of codon aliases (=1/20).

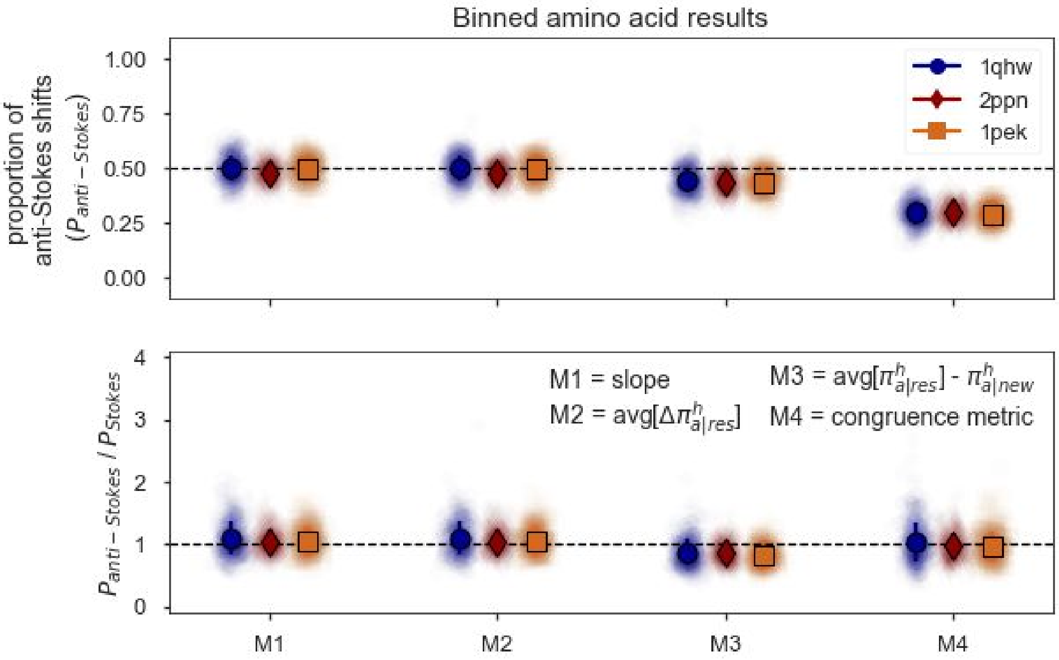


**Figure S2.** Proportion of evolutionary anti-Stokes shifts considering shifts in propensities in bins of amino acids that tend to interchange rapidly and that tend to have similar chemical properties. Amino acids were grouped as: AST, C, DE, FY, GN, HQ, IV, KR, LM, P, W. Evolutionary shifts were calculated based on the sum of propensities for all amino acids in a specific bin. (A) Approximately half of substitutions are followed by evolutionary anti-Stokes shifts based on metrics M1-3, a slightly lower proportion (approx. 0.3) was estimated under M4. (B) Across all metrics M1-4, evolutionary Stokes and anti-Stokes shifts occur at similar frequencies ($P_{anti-Stokes} / P_{Stokes} \approx 1$).

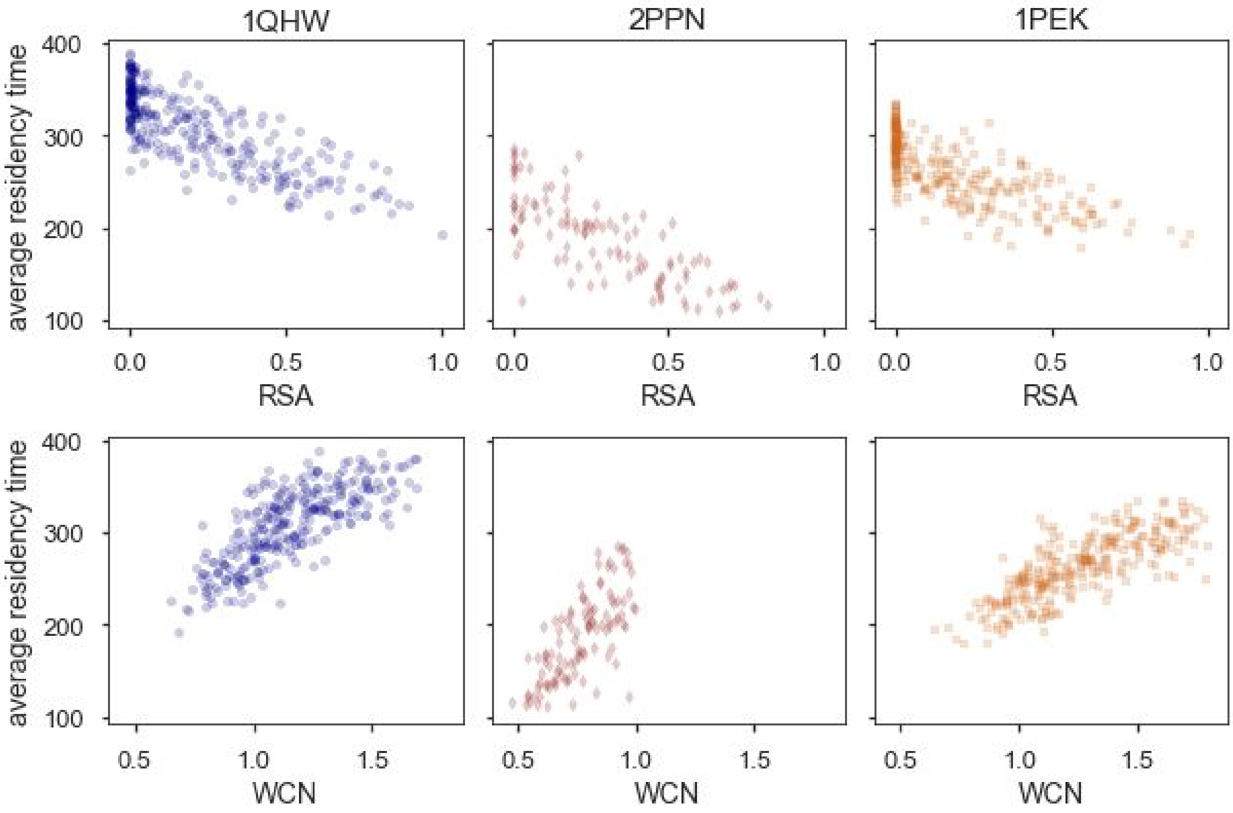


**Figure S3.** Relationship between average amino acid residency time and relative solvent accessibility (RSA, top row), and weighted contact number (WCN, bottom row) for three proteins (1qhw, 2ppn, 1pek).

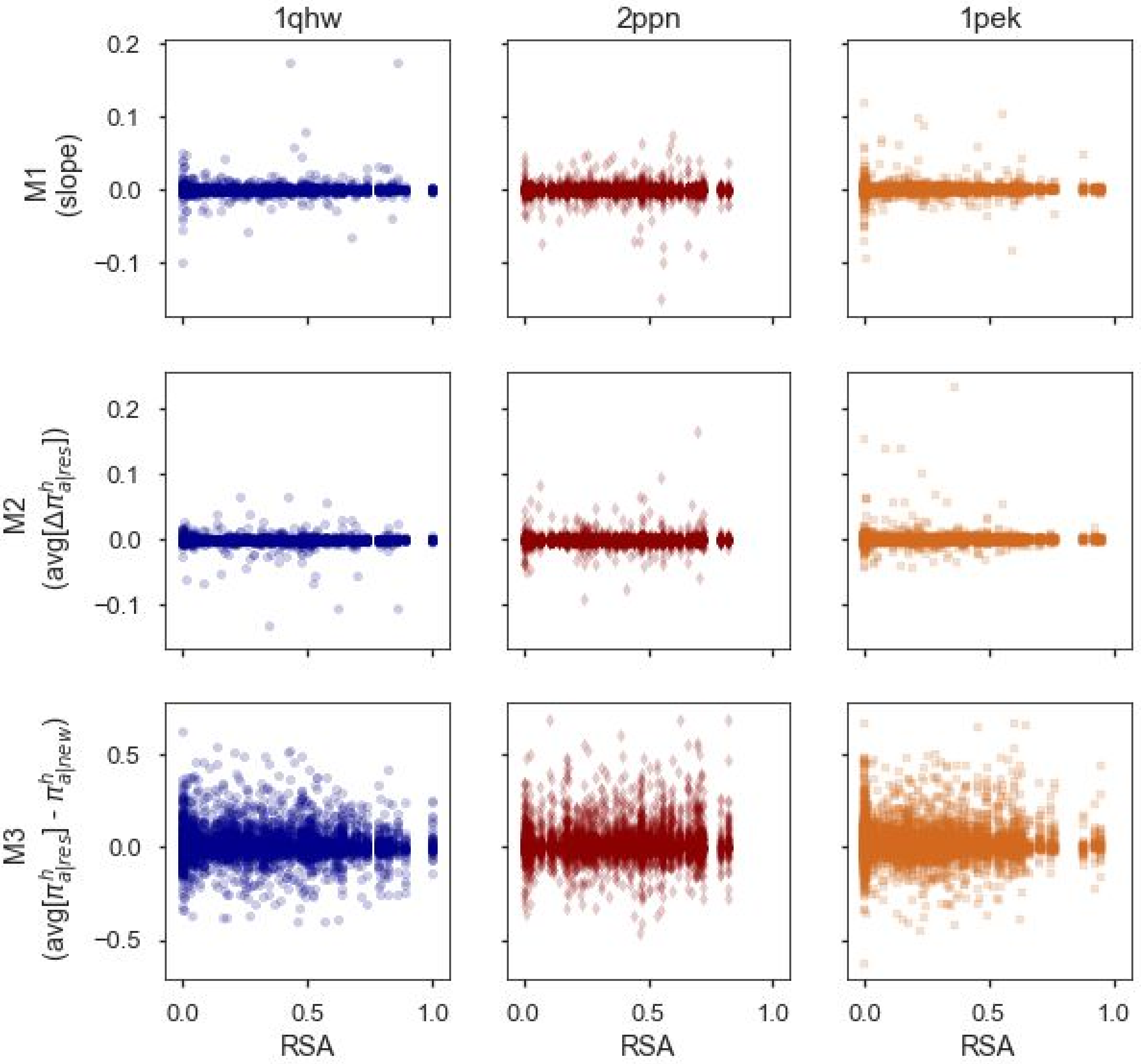


**Figure S4.** Relationship between quantitative metrics (M1, M2, and M3) and relative solvent accessibility (RSA).

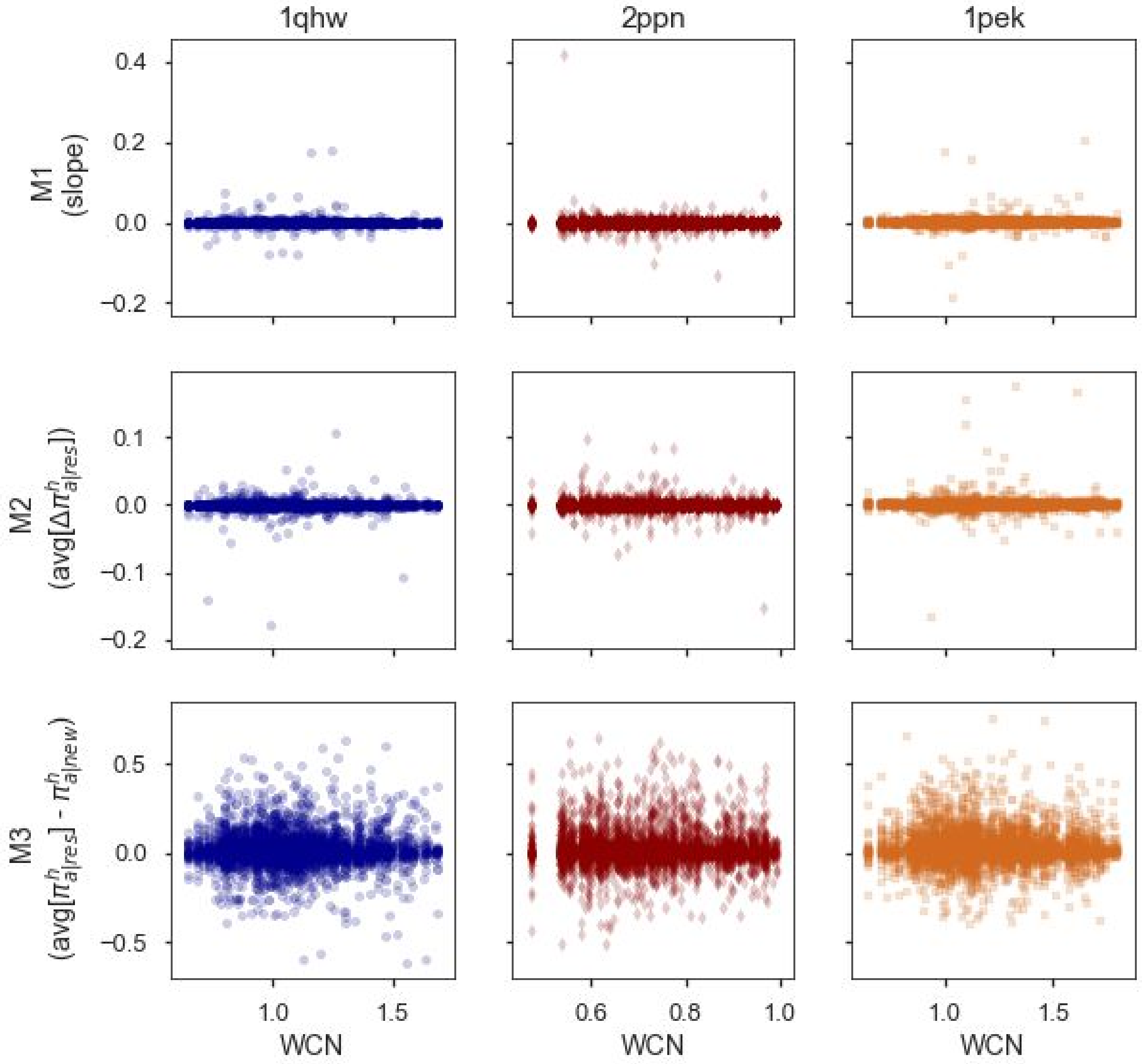


**Figure S5.** Relationship between quantitative metrics (M1, M2, and M3) and weighted contact number (WCN).

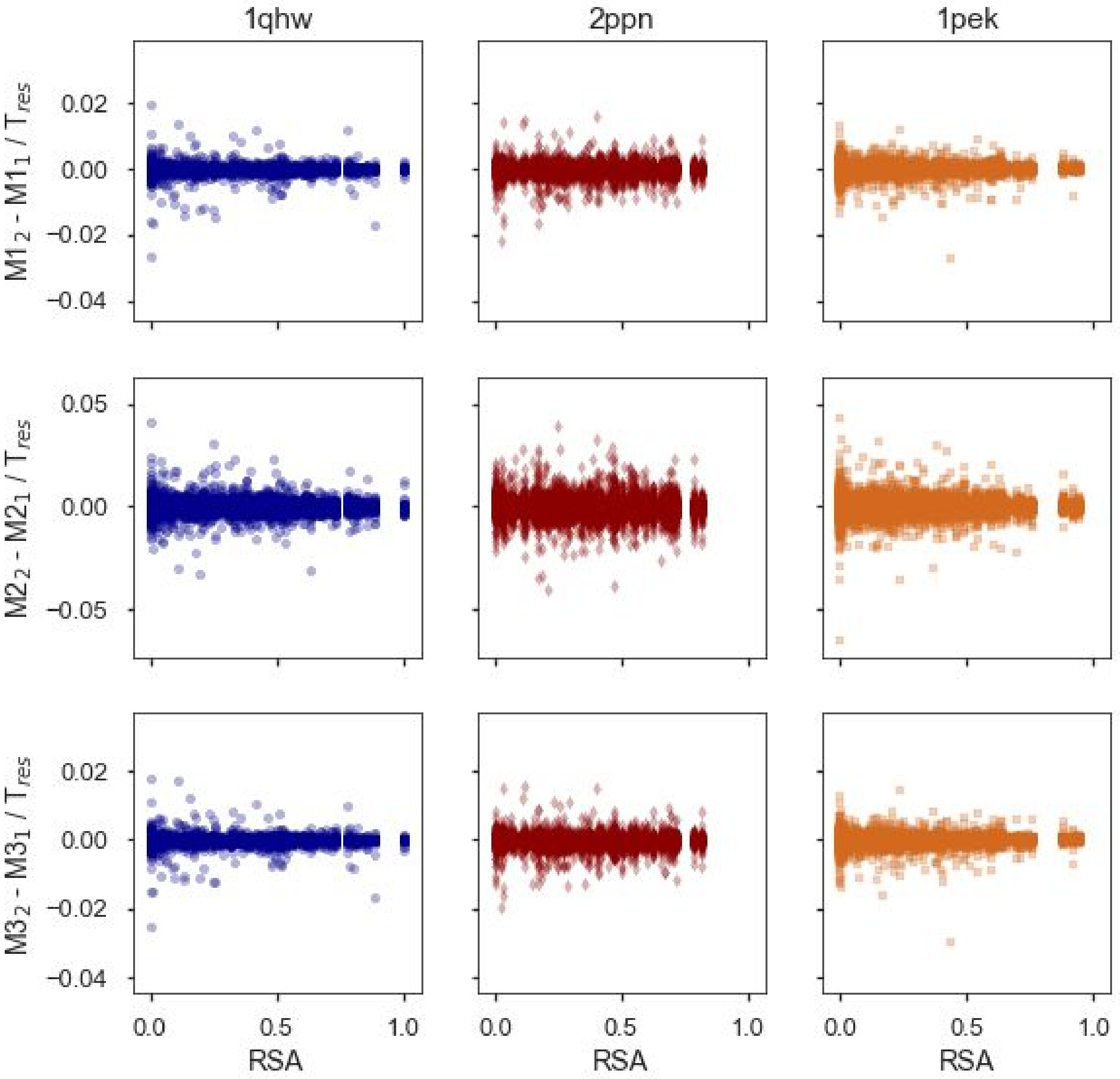


**Figure S6.** Relationship between the rate of acceleration (or deceleration) of evolutionary shifts calculated based on metrics M1, M2, and M3 and relative solvent accessibility (RSA).

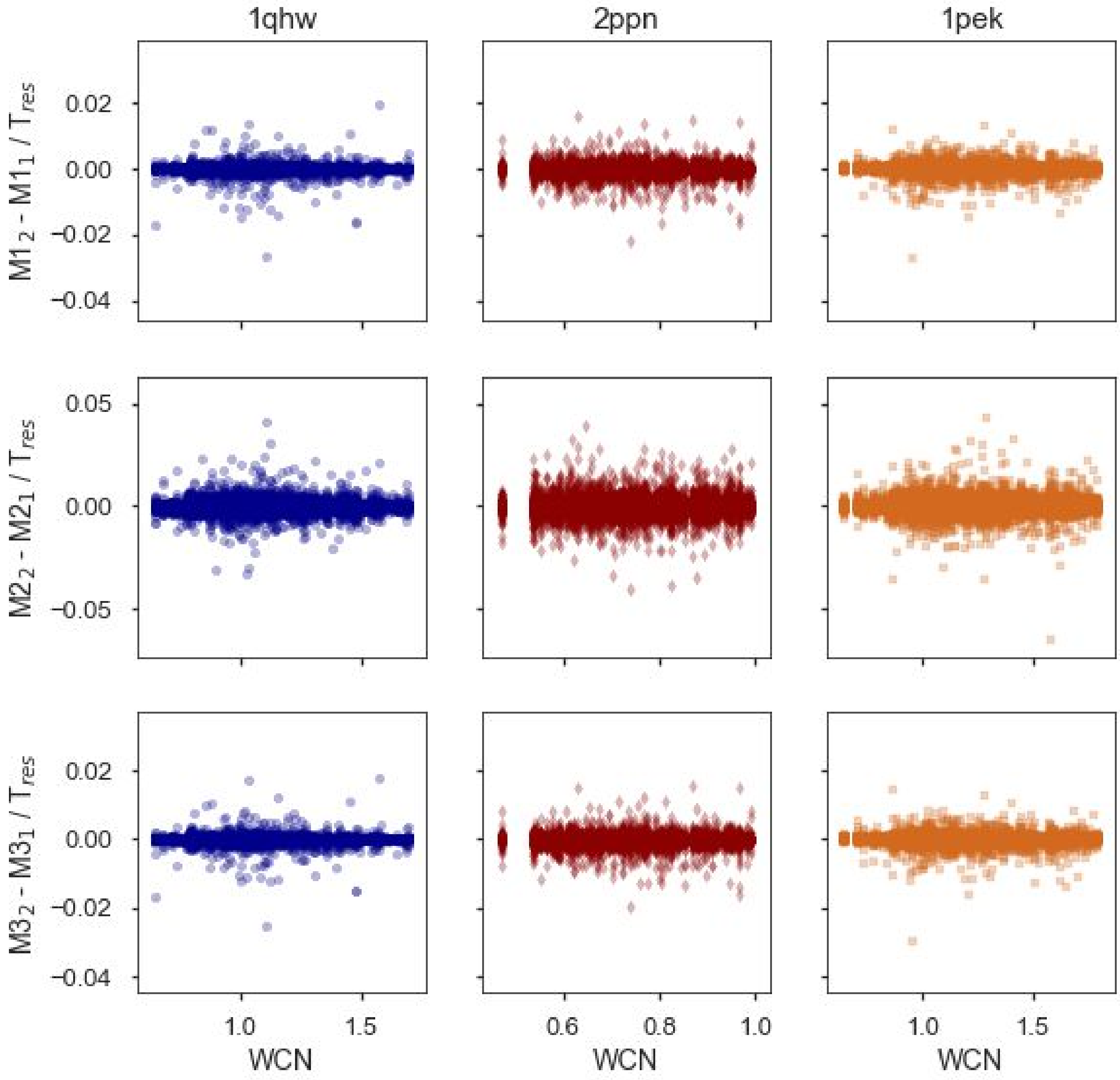


**Figure S7.** Relationship between the rate of acceleration (or deceleration) of evolutionary shifts calculated based on metrics M1, M2, and M3 and weighted contact number (WCN).

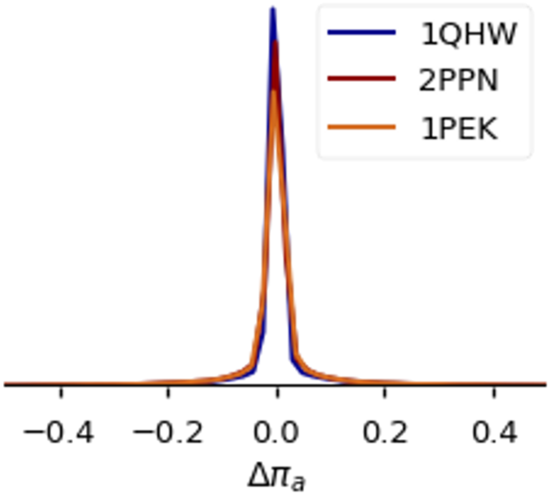


**Figure S8.** Distributions of the observed changes in resident amino acid propensities ( $\Delta\pi_a$ ) throughout the simulations of three protein structures: 1qhw, 1pek, and 2ppn.

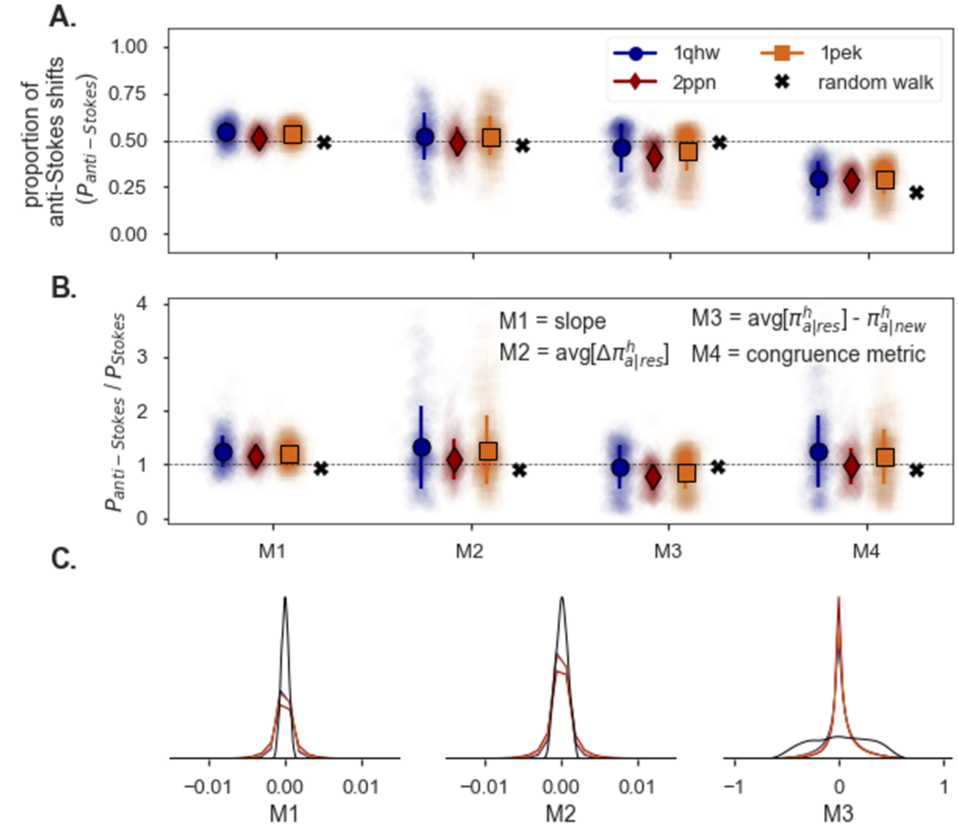


**Figure S9.** Evolutionary Stokes and anti-Stokes shifts occur with similar frequencies under a bounded random walk model (between 0 and 1) and the stability-constrained model. (A) The proportion of evolutionary anti-Stokes shifts as estimated based on metrics M1-4 from simulations of a bounded random walk and stability-constrained simulations with protein structures 1qhw, 2ppn, and 1pek. (B) The relative frequency of evolutionary anti-Stokes shifts compared to evolutionary Stokes shifts ($P_{anti-Stokes} / P_{Stokes}$). (C) Distributions of metrics M1-3. All metric values were symmetric and centered at zero.

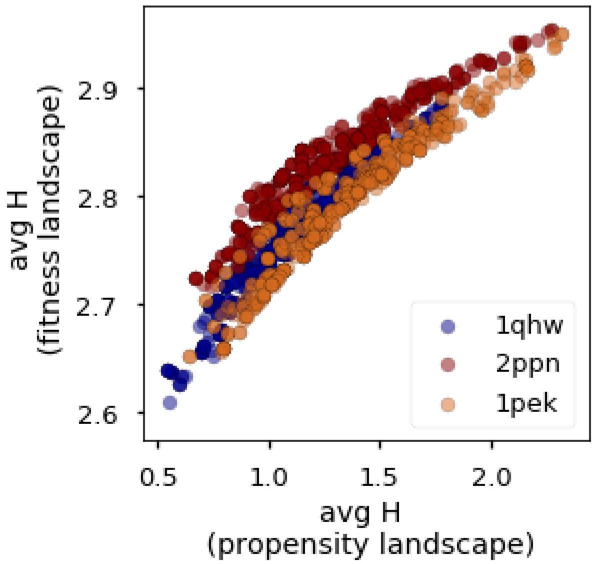


**Figure S10.** Relationship between the Shannon entropy for propensity versus fitness landscapes. Reported are the average entropy values over all sites given a particular background sequence from a single simulation trial for each of three protein structures (1qhw, 2ppn, and 1pek).

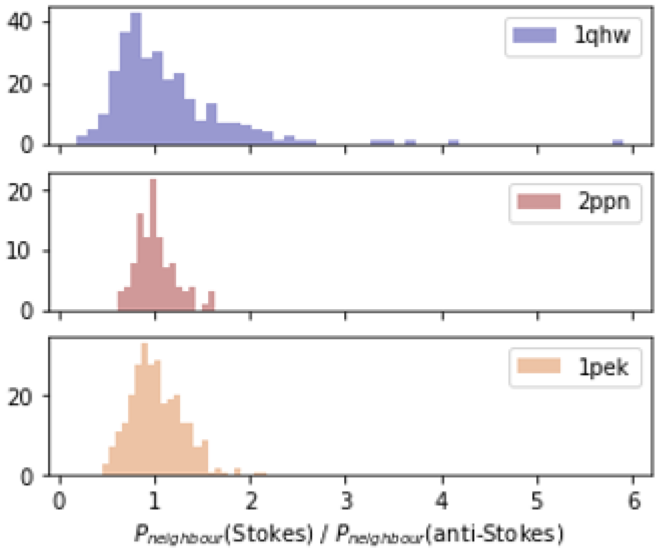


**Figure S11.** Distributions of the relative proportion of neighbouring nonsynonymous substitutions when an evolutionary Stokes shift occured ($P_{neighbour}$(Stokes)) compared to when anti-Stokes shifts occurred ($P_{neighbour}$(Stokes)). Evolutionary Stokes and anti-Stokes shifts were classified based on metric M4. Sites are considered to be neighbouring if their beta carbons are within 7 Angstroms (if the amino acid is Glycine distance is calculated relative to the alpha carbon).

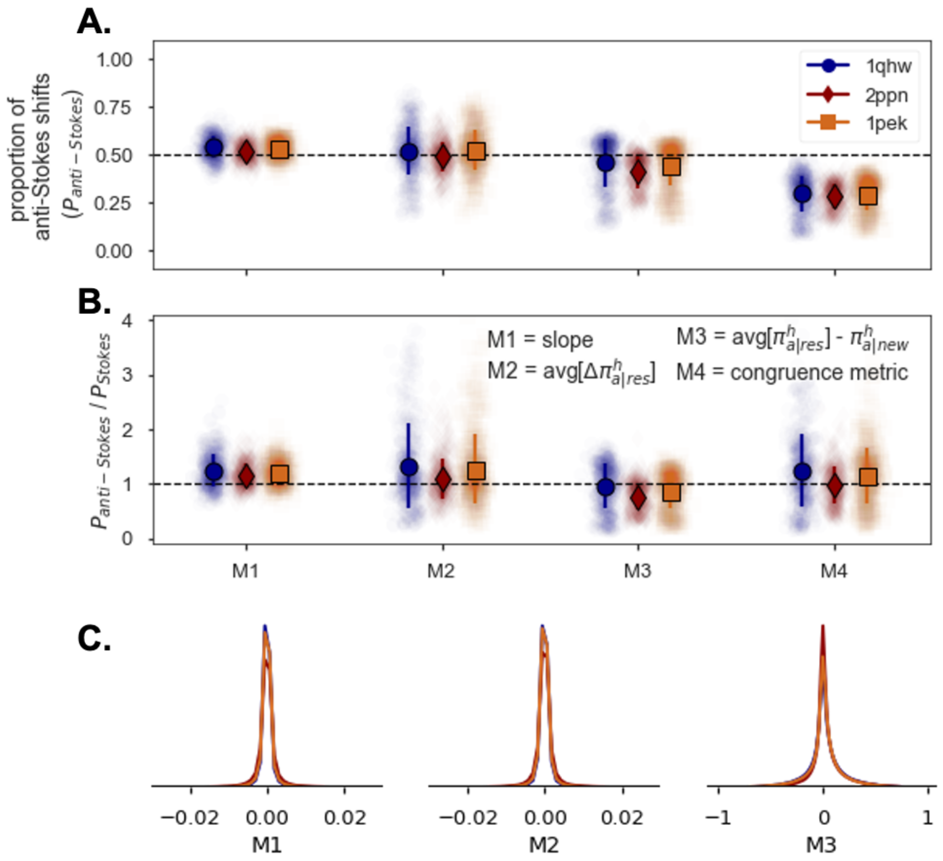


**Figure S12.** Proportions of evolutionary Stokes and anti-Stokes shifts calculated including partial windows, where for example an amino acid is accepted but the simulation ends prior to its replacement. (A) Approximately half of substitutions are followed by evolutionary anti-Stokes shifts based on metrics M1-3, and approx. 0.3 under M4. (B) Across all metrics M1-4, evolutionary Stokes and anti-Stokes shifts occur at similar frequencies ($P_{anti-Stokes} / P_{Stokes} \approx 1$). (C) The values of M1-3 are normally distributed and centered at zero which suggests that both the magnitude and frequencies of Stokes and anti-Stokes shifts are balanced.
